# High-resolution model of Arabidopsis Photosystem II reveals the consequences of digitonin-extraction

**DOI:** 10.1101/2021.01.15.426676

**Authors:** André T. Graça, Michael Hall, Karina Persson, Wolfgang P. Schröder

## Abstract

In higher plants, the photosynthetic process is performed and regulated by Photosystem II (PSII). *Arabidopsis thaliana* was the first higher plant with a fully sequenced genome, conferring it the status of a model organism; nonetheless, a high-resolution structure of its Photosystem II is missing. We present the first Cryo-EM high-resolution structure of Arabidopsis PSII supercomplex with average resolution of 2.79 Å, an important model for future PSII studies. The digitonin extracted PSII complexes demonstrate the importance of: the LHG2630-lipid-headgroup in the trimerization of the light-harvesting complex II; the stabilization of the PsbJ subunit and the CP43-loop E by DGD520-lipid; the choice of detergent for the integrity of membrane protein complexes. Furthermore, our data shows at the anticipated Mn_4_CaO_5_-site a single metal ion density as a reminiscent early stage of Photosystem II photoactivation.

## Introduction

Photosynthesis is the most important natural process existing on our planet, in which solar radiation is converted into biomass^1,2^. It provides food, oxygen to breathe and renewable, sustainable energy. It is also one of the few processes that reduce amounts of greenhouse gas carbon dioxide in the air^3^. The weed plant *Arabidopsis thaliana* (thale cress or mouse-ear cress) became a model organism for plant research due to several traits such as size, generation time, accessibility, manipulation, and genetics^4^. The use of Arabidopsis as a model organism was enhanced as its full genome was sequenced and published in the year 2000^5^.

Photosystem II (PSII)—a key player in photosynthesis—is a multi-subunit light-driven water:plastoquinone oxidoreductase^6^ located in the thylakoid membrane of chloroplasts^7^. Functional PSII has a dimeric core, surrounded by light-harvesting complexes (LHCII), and thus constitutes one of the largest protein complexes in nature with more than 60 protein subunits and a molecular mass exceeding 1400 kDa^8^. In its monomeric composition, Photosystem II reaction centre is built of the essential D1-D2 heterodimer, that hosts several cofactors responsible for the light-induced charge separation and water oxidation^6,8^. These core proteins are connected to an internal core chlorophyll *a* binding antenna, composed of CP43 and CP47 proteins^9^. Chlorophyll *a/b* binding minor antenna proteins (CP24, CP26, CP29) link the major chlorophyll *a/b* binding LHCII to the PSII core^10,11^. The trimeric LHCII is the most abundant chlorophyll-binding protein assembly in eukaryotic photosynthetic organisms, and it is constituted of three polypeptides, Lhcb1Lhcb1-3^12,13^. Several of these trimeric assemblies can bind simultaneously to a PSII core complex at specific locations. Their relative location to the PSII dimeric core (C_2_) and the strength of the PSII-LHCII interaction creates4 distinct classes^14^: strong (S), medium (M), loose (L) and naked (N). The N-LHCII type, recently discovered in the green algae *Chlamydomonas reinhardtii*^15–17^; and the less characterized L-LHCII type can be found in land plants^18^, although trimers are not abundantly associated with PSII under optimal light conditions.

The lumen side of land plants PSII complex harbours four extrinsic proteins (PsbO, PsbP, PsbQ and PsbTn). Additionally, there are at least 10 low molecular mass proteins (< 10 kDa) at various sites, between major proteins and in the outskirts of the complex^19^. An immense network of pigments, lipids and other essential cofactors are known to bind to the different proteins of the complex^8,20–22^. The pigments—chlorophylls and carotenoids—are essential for light absorption while the lipids main function is the ligation of different subunits at several oligomerization interfaces and the binding of some pigments^23^.

Structural studies of Photosystem II were for an extended period restricted to X-ray crystallography methods which demand high concentrations of pure homogeneous and stable complexes and the capability of symmetric assembly to form crystals^24,25^. Thus, structural studies of bacterial PSII from *Thermosynechococcus elongates* and *Thermosynechococcus vulcanus* have been commonly performed due to their high stability and accessible large batch extraction. Such attempts resulted in several high-to near atomic resolution structures, between 1.9-3.8 Å average resolution^21,26–29^. The bacterial PSIIcc is expected to be very similar to the dimeric PSII core of higher plants (PSII C_2_), however there are several distinct differences^30^: i. bacterial PSII lacks chlorophyll *b*; ii. the light-harvesting antenna is integrated into the membrane in higher plants, while it is peripheral in bacteria (phycobilisomes); iii. the extrinsic proteins differ between bacteria and higher plants; iv. the pigments and the lipids not always share conserved binding-sites between species of different kingdoms.

Higher plant PSII supercomplexes have traditionally been extracted from thylakoid membranes using dodecylmaltoside (α- or β-isoforms) and digitonin detergents^31–33^. Other detergents have been screened for the purpose, but results show that PSII complexes do not remain stable in a wide variety of detergents^31^. The high concentration of detergents, and the different sizes of PSII complexes extracted from the thylakoid membrane, are two factors that hindered the formation of stable crystal assemblies^34,35^. A new opportunity to study the higher plants PSII structure came with the recent developments of single-particle cryo-electron microscopy (EM)^36^. This method allows acquisition of structural information from large protein complexes at resolutions similar to X-ray crystallography. As the sample is plunge frozen and the imaged particles are segregated into classes upon computational analyses, the sample does not need to be as pure and enriched as for crystallisation and their level of stability is only relevant up to the moment of plunge freezing^36^. The first higher plant PSII Cryo-EM structures were recently reported for; *Spinacia oleracea* (Spinach) at 3.2 Å resolution (PDB: 3JCU)^22^; *Pisum Sativum* (Pea), with a resolution of 2.7 Å and 3.2 Å (PDB: 5XNL and 5XNM, respectively)^20^, the former consisting of a smaller C_2_S_2_ complex. However, concerning *Arabidopsis thaliana*, the best structural model of PSII (PDB: 5MDX)^33^—extracted from a mutant phenotype—has a resolution of 5.3 Å.

Photosystem II oxygen production concurrently exposes the complex to oxidative stress, leading to damage of the D1 protein^37^. To cope with this, plants have developed various protective mechanisms, such as the ability to repair the PSII complex by replacing the D1 protein^38^. Under high-light conditions, i.e. when plants absorb more light than they can use for photosynthesis the D1 protein is replaced every 20-30 minutes, which means that the assembly state of PSII is in a constant switch between functional and repair/inactive state. To become functional, PSII needs to correctly assemble the site for water oxidation; a process driven by light and known as photoactivation^37,39^. At distinct steps of photoactivation, manganese ions (Mn^2+^) interact with specific residues of the D1 protein, while being oxidized^39^. One calcium ion (Ca^2+^) and five oxygen atoms bridging the metal ions are indispensable to form the functional high-valence manganese cluster, Mn_4_CaO_5_. When the full water oxidation complex is formed, Mn_4_CaO_5_ is coordinated to several D1 and CP43 amino acids^26^. Even though most of the photoactivation steps are still to be understood, there is evidence that manganese and calcium ions compete to bind at the <u>h</u>igh-<u>a</u>ffinity Mn-binding <u>s</u>ite (HAS)^39^—involving at least the D1 residue Asp170—in an early step of photoactivation. The following steps are the oxidation of the first-bound Mn^2+^ and the so-called ‘dark rearrangement’, proposed as a light-independent dynamic where the C-terminal of the D1 protein adopts a new structural conformation like the observed in the Mn_4_CaO_5_ binding pocket of mature PSII^39^. In an attempt to gain new information of this activation process, Gisriel and colleagues^40^ isolated monomeric PSII from *Synechocystis* sp. PCC 6803 that were EDTA-washed and buffer exchanged. This recent study showed that the Mn_4_CaO_5_-binding pocket undergoes major rearrangements upon assembly of the OEC, before the formation of a fully functional PSII.

Here we present the first high-resolution structure of Arabidopsis PSII C_2_S_2_M_2_ supercomplex. This model shows that the absence of the extrinsic PsbP and PsbQ proteins leads to a disordered C-terminus of the reaction centre D1 protein which we speculate to be important for the assembly process of the manganese water splitting complex of PSII. Furthermore, new knowledge of protein-protein, protein-ligand and ligand-ligand interactions is attained by accessing the consequences of the digitonin molecules bound to PSII.

## Results

### The overall structure of the Arabidopsis Photosystem II complex

The imaged Photosystem II C_2_S_2_M_2_-type complexes were isolated from wild-type Arabidopsis BBY membrane fragments^41^ solubilised with a mixture of digitonin and β-DDM detergents. Visual inspection of the obtained micrographs suggested a high level of sample homogeneity. At the same time, the implemented *ab initio* 3D reconstruction strategy allowed us to further filter the dataset to a more homogeneous subset of 100 712 PSII particles. The 3D reconstruction, with imposed C2 symmetry, yielded a map at an overall resolution of 2.79 Å for the C_2_S_2_ supercomplex, according to FSC_0.143_ golden standard (Appendix, Fig S1 and Table S1). However, the resolution was found to be heterogeneous, with a higher resolution of 2.5 Å throughout the core region and a lower resolution of 6.0 Å in the outer regions of the complex (Appendix, Fig S2). An additional data collection using a Volta phase plate was performed on the same grid, yielding a map at 3.6 Å global resolution (C_2_S_2_M_2_) that helped to resolve certain peripheric regions of the complex such as additional amino acids of stromal terminal regions of certain subunits (e.g.: N-terminal of the D1 protein).

The Arabidopsis PSII C_2_S_2_M_2_ supercomplex has a two-fold symmetry structure (Appendix, Fig S3), with dimensions estimated to be 280 x 180 x 107 Å (LWH). Our model revealed the presence of 17 different protein subunits in the PSII core complex (Appendix, Table S2), including PsbA (D1), PsbB (CP47), PsbC (CP43), PsbD (D2), PsbE, PsbF, PsbH, PsbI, PsbK PsbL, PsbM, PsbO, PsbTc, PsbTn, PsbW, PsbX and PsbZ. An overview of the modelled proteins and their location in the Arabidopsis PSII monomer is shown in **Fig 1**. Comparing this model to the Pea and Spinach models, the spatial distribution of the core proteins did not reveal any dramatic change, which suggests that the PSII core is structurally and compositionally very well conserved among higher plants. The light-harvesting proteins CP24, CP26, CP29 and four LHCII trimers were identified and modelled (Appendix, Table S3). The highly resolved features of the map, especially of the core complex proteins, allowed that several new structural observations could be made including: 35 N-terminal amino acids of the D1 protein, lacking on the previous medium-resolution Arabidopsis PSII model (PDB: 5MDX), were fully modelled; similarly, N- and C-termini of D2 were modelled to a greater extent when compared to 5MDX; the nuclear-encoded extrinsic protein PsbTn; per PSII monomer, one chloride (Cl^-^) ion was found bound to the complex (Appendix, Fig S4) in the vicinity of the OEC; the vast carotenoid pigment network and the lipids present in PSII; and 141 water molecules were modelled.

**Figure 1.**
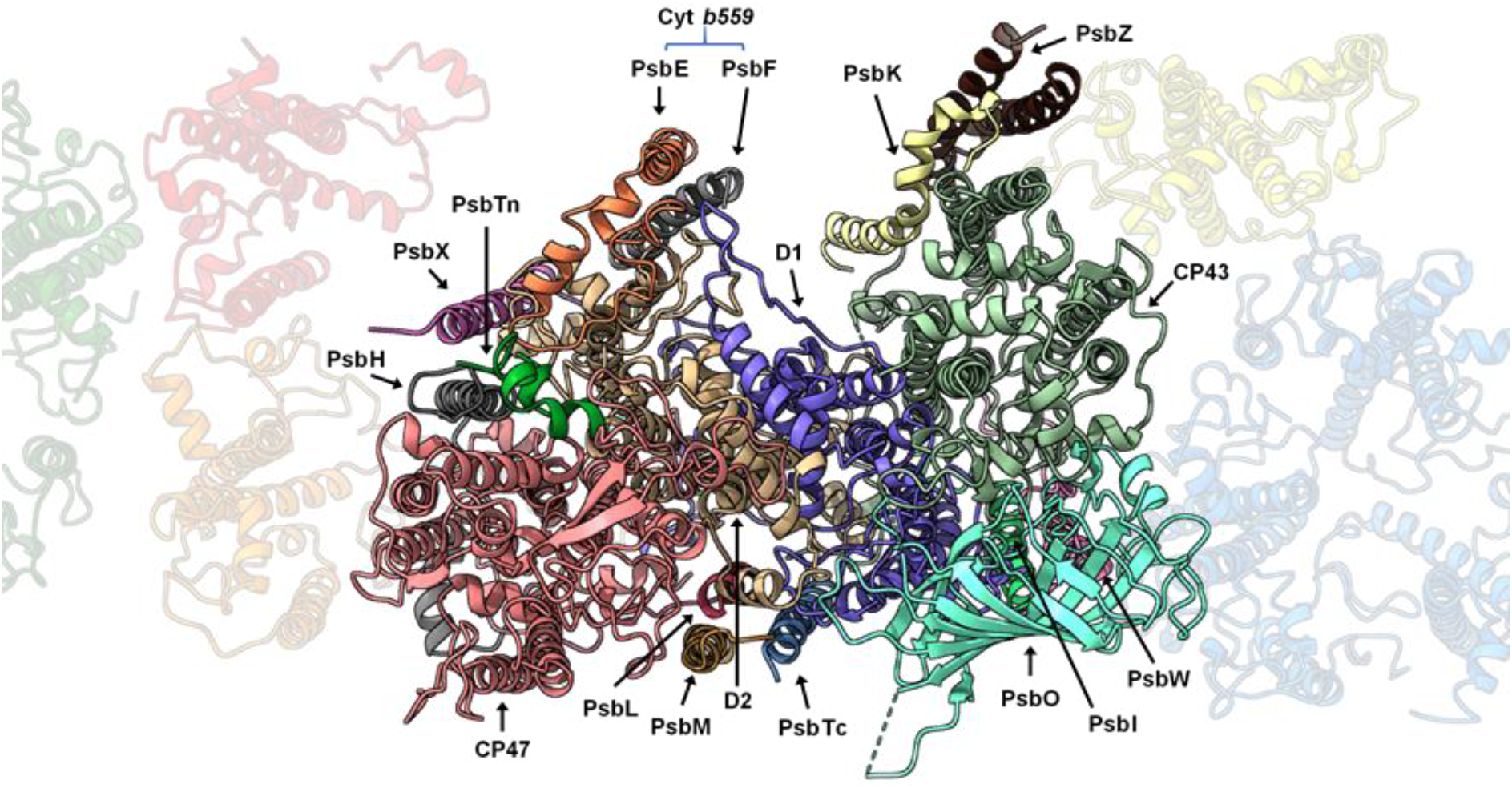
Localization of the Photosystem II core proteins. An entire PSII monomer is displayed in first plane, as seen from the lumen side of the thylakoid membrane, with the peripheral antennae in the background.

The main four core proteins—D1, D2, CP47 and CP43—bind together 73 cofactors of which 35 are chlorophylls *a* (Appendix, Fig S5). The current resolution enabled us to model the correct spatial orientation of chlorophylls’ chlorin rings and often the full extension of their hydrophobic tails. Unobserved in the previous medium-resolution map of Arabidopsis PSII, a density for CP29 chlorophyll *a* at position 616 is clearly visible in our EM map.

In total, 162 chlorophyll *a*, 60 chlorophyll *b*, four pheophytin *a*, and two heme molecules were modelled. Additionally, 60 lipids (**Fig 2A**), 42 carotenoids (Appendix, Fig S6), and two bicarbonate molecules were visible in our EM map and for the first time described in an Arabidopsis PSII model (Appendix, Table S4). Besides the cofactors mentioned above, 22 digitonin molecules were identified and modelled (**Fig 2B**). The well-defined densities of these detergent molecules and those of the several PSII cofactors present in our model, demonstrate the quality of the obtained EM map (Appendix, Fig S7).

**Figure 2.**
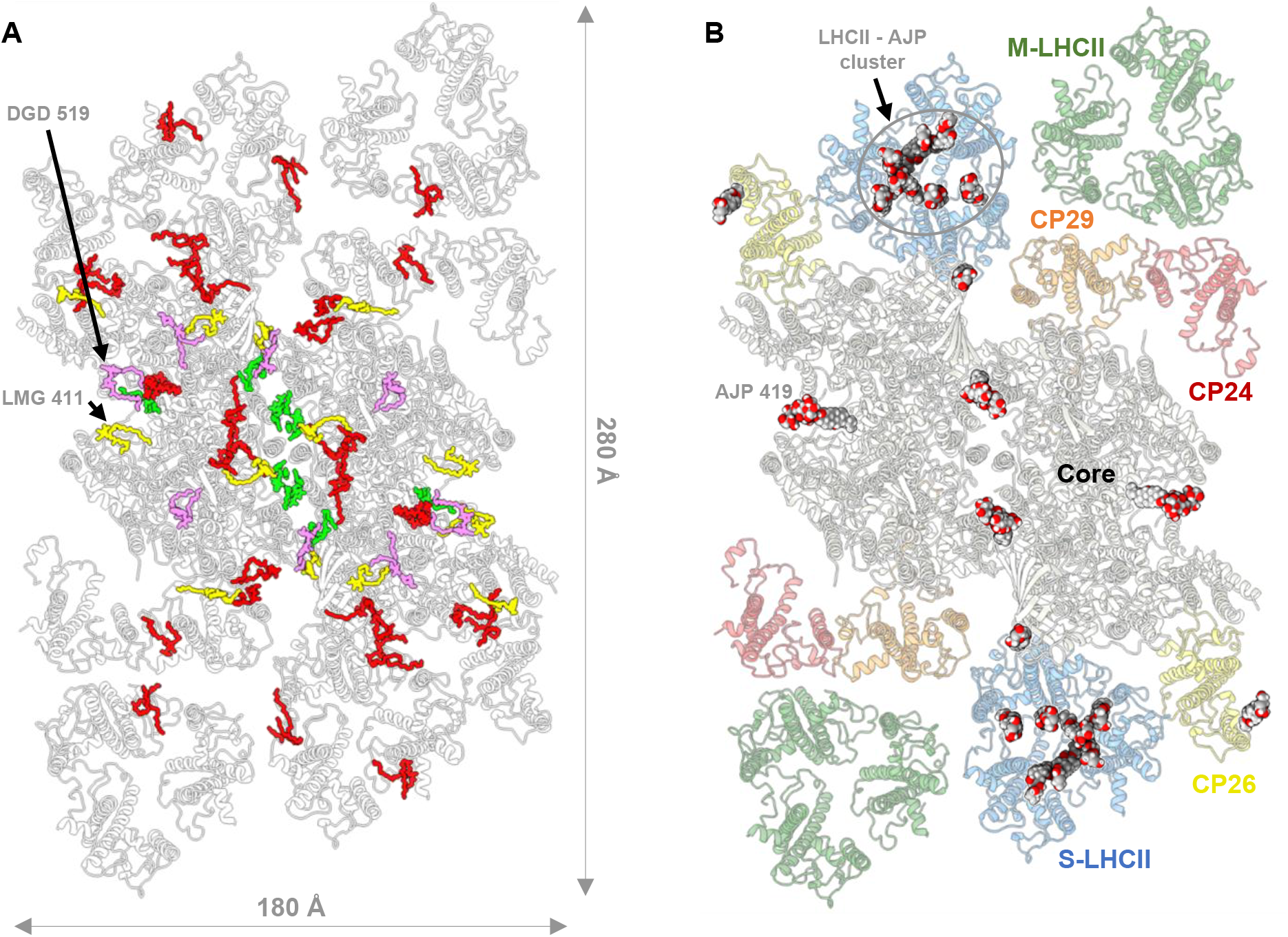
**A**, Distribution of the lipids in Arabidopsis PSII, as seen from the lumenal side: monogalactosyldiacylglycerol (MGDG/LMG, in yellow), digalactosyldiacylglycerol (DGDG/DGD, in violet), sulfoquinovosyldiacylglycerol (SQDG/SQD, in green) and phosphatidylglycerol (PG/LHG, in red). **B**, Distribution of the modelled digitonin molecules (ligand code: AJP) in our PSII supercomplex, as seen from the lumenal side. The secondary structure of the protein backbones is represented in cartoon style and the detergent molecules represented by spherical atomic representation.

Of the four extrinsic proteins (PsbO, PsbP, PsbQ, and PsbTn), PsbO and PsbTn were detected in our EM map. The Arabidopsis genome encodes two isoforms of the PsbO protein, referred to as PsbO1 and PsbO2, that share 90% sequence identity. PsbO1 is expressed at a higher level than its counterpart and it is suggested to be bound to PSII and involved in its activity^42^. With a high-resolution structure of PSII we can differentiate between some of the non-conserved residues between the two isoforms. The densities corresponding to PsbO residues at positions 128 and 187, fit better to a proline and an asparagine as in PsbO1, than the corresponding alanine and lysine residues of PsbO2. Therefore, we have modelled PsbO in accordance with PsbO1 sequence (Appendix, Table S2). Nevertheless, the local resolution of our EM corresponding to PsbO is lower than the PSII core, suggesting that the entire protein has some flexibility or that a lesser percentage of PSII supercomplexes may indeed bind the PsbO2 isoform.

The analysis of individual 2D and 3D classes, the detection of PsbTn with low occupancy, and the absence of PsbP, PsbQ and the neighbouring transmembrane protein PsbR, suggest that the binding of these proteins is highly sensitive to protocol variations. PsbP, PsbQ and PsbR subunits were also reported missing in the previous Arabidopsis PSII model and in the unstacked model of Pea PSII. The absence of these proteins is probably owing to thylakoid membrane solubilizations performed at pH 7.5 using high concentrations of α-DDM or lower concentrations of DDM with additional digitonin detergents (Appendix, Table S5). We do not disregard that the strength of the binding of PsbP and PsbQ to the PSII complex may differ between land plants. This argument is supported by the presence of these subunits (in substoichiometric amount) in the Pea PSII electrostatic potential map, extracted at pH 7.5^22^.

In contrast to the previous medium-resolution Arabidopsis PSII cryo-EM structure from PsbS knock-out mutant^33^, the hereby presented atomic model is the first PSII structure derived from wild-type phenotype of *Arabidopsis thaliana*. Although the PsbS protein was present in the starting BBY sample, we were not able to detect any density corresponding to this protein in any of the investigated 2D projections or the final EM map. Thus, the location of the enigmatic energy quenching^43^ PsbS protein remains to be determined.

Of the major antennae complexes, we identify two strongly bound LHCII (S-LHCII) and two moderately bound (M-LHCII). S-LHCII contains three monomers A, B and C (S-LHCII_A_, S-LHCII_B_ and S-LHCII_C_); these are within exciton energy transfer (EET) distance (∼20Å) to other PSII subunits at four different places, S-LHCII_B_ to CP26, S-LHCII_A_ to CP43 and CP29, and S-LHCII_C_ to the adjacent M-LHCII_1_ (Appendix, Fig S5 and Table S6), which probably explains the strongly bound features of this LHCII subunit.

In general, the region around M-LHCII trimers and CP24 could not be easily resolved due to the: higher flexibility of this region in relation to the core of the complex; diminished occupancy arising from the heterogeneity of the sample with an unquantifiable number of C_2_S_2_ particles present in our dataset; destabilisation caused by the used detergent mix. The low resolution of the M-LHCII-CP24 region, did not allow to determine which of the *lhcb*1, *lhcb*2 or *lhcb*3 gene product^44^ correspond to which LHCII monomer. M-LHCII trimers— monomers modelled according to Lhcb3 (in contact with CP24) and two Lhcb1.4 protein sequences—and associated CP24 protein were placed into their respective densities. Besides their relative location to the core, detailed structural information cannot be inferred from the current EM map thus, ligands were only modelled for unequivocal densities. We note that divergences at the sequence level may occur since our EM map does not present resolved side-chain densities for the M-trimer Lhcb proteins.

### The peripheral low molecular mass proteins of the PSII core

Three low molecular mass proteins, PsbW, PsbH and PsbZ, were found located in-between the PSII core and the antenna complexes (**Fig 1**). The PsbW protein links the LHCII monomer A to CP43^45^; meanwhile, the PsbH is bridging the CP29 to CP47, and the PsbZ bridging the CP26 to the CP43 protein. As none of them has been shown to bind any pigment, likely, they are not directly involved in energy transfer from the antenna to the reaction centre. Thus, suggesting that their primary function is to support the association of the antennae to the core of the PSII complex.

The well-resolved side chains densities for the PsbW protein enabled us to fully fit the N-terminal tail and model its remaining residues; and address the unknown structure of its C-terminus (**Fig 3**). We concluded that the N-terminal tail was misplaced in the previous medium-resolution PSII model and identified the amino acids PsbW-Asn99, PsbW-Trp103, and LHC_A_-Asn122 (Appendix, Fig S8), as being the potential motifs that anchor the S-LHCII trimer to PsbW through electrostatic interactions.

**Figure 3.**
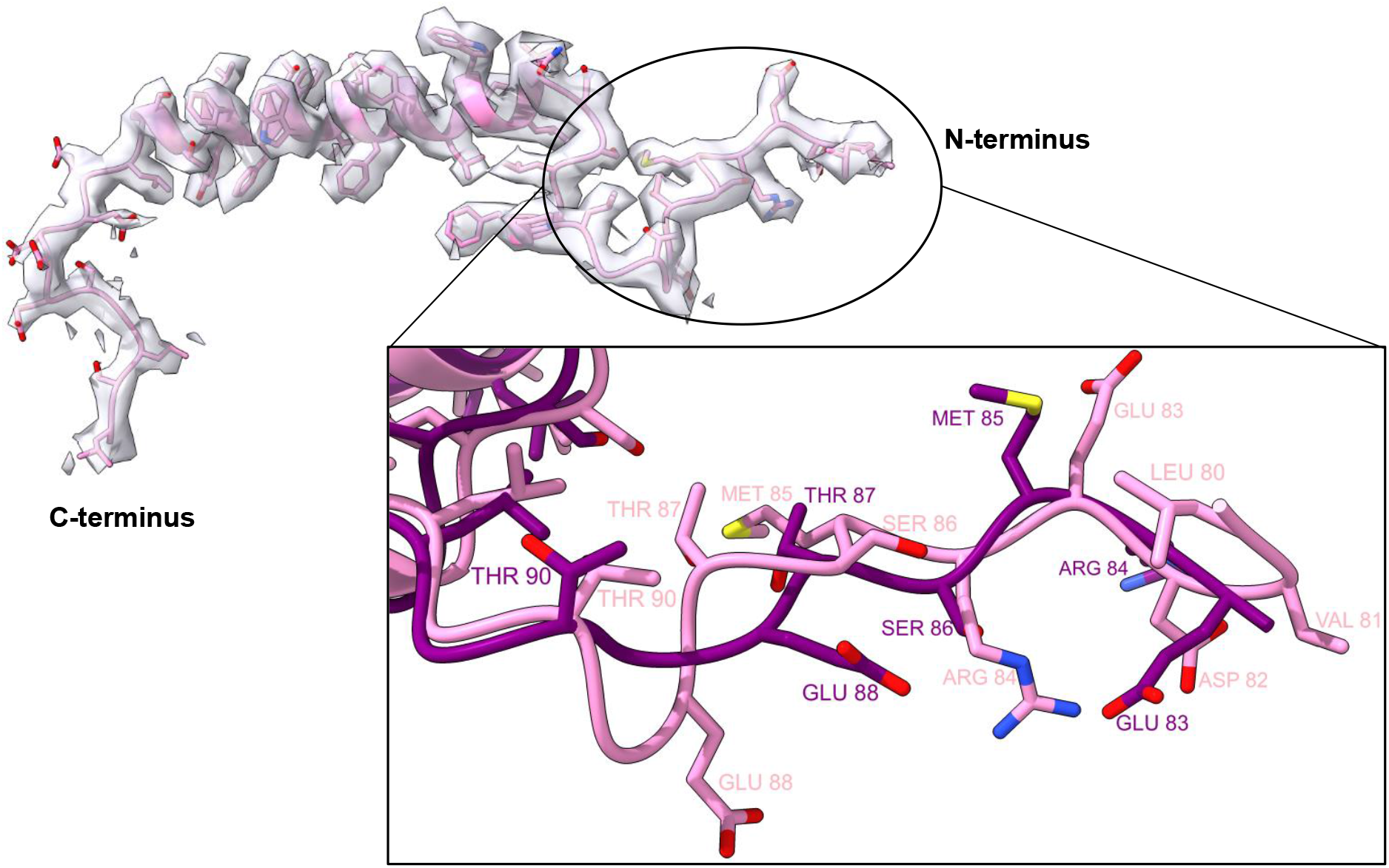
The low molecular weight protein PsbW fit into its respective density. Inset: Dissimilar modelling of the N-terminus of PsbW between #??? and 5MDX (purple) Arabidopsis PSII models

PsbW is suggested to interact with the PsbH subunit of the opposite PSII monomer and in this way participate in the stabilisation of the PSII dimer^46^. This interaction of electrostatic nature is supported by our finding that the negatively charged C-terminal tail of the PsbW protein and the positively charged N-terminal tail of PsbH are both on the stromal side close to each other. Phosphorylation of PsbH would in this case hinder such interaction, triggering the disassembly of PSII and preparing it for repair^47^. Although the C-terminal tail of PsbW adopts a conformation that follows in the opposite direction of PsbH, this possibility cannot be excluded since no cryo-EM map of higher plant PSII model was able to resolve the 12 amino acids of PsbH N-terminal tail.

We used the sequence motif PSLK (Pro81-Lys84) of the PsbX protein and the corresponding side chains densities, to correctly place the subunit and resolve four additional amino acids of its N-terminus (Appendix, Fig S9). Our placement of PsbX is shifted four to five amino acids (≈ 7.9 Å, alpha-carbon shift) towards the stromal side compared to the previous medium-resolution Arabidopsis PSII model.

### Digitonin impact several oligomerization interfaces between PSII proteins

The superposition of Pea and Spinach PSII structures to our PSII model reveals that the distance between the two PSII monomeric cores is longer in Arabidopsis PSII. This is probably explained due to the presence of two digitonin molecules at the monomer-monomer interface, affecting the intermonomer distance (**Fig 4** and Appendix, Fig S10-11). However, the stability of the dimer does not seem to be affected as the purification of a homogenous sample of dimerised PSII was possible.

**Figure 4.**
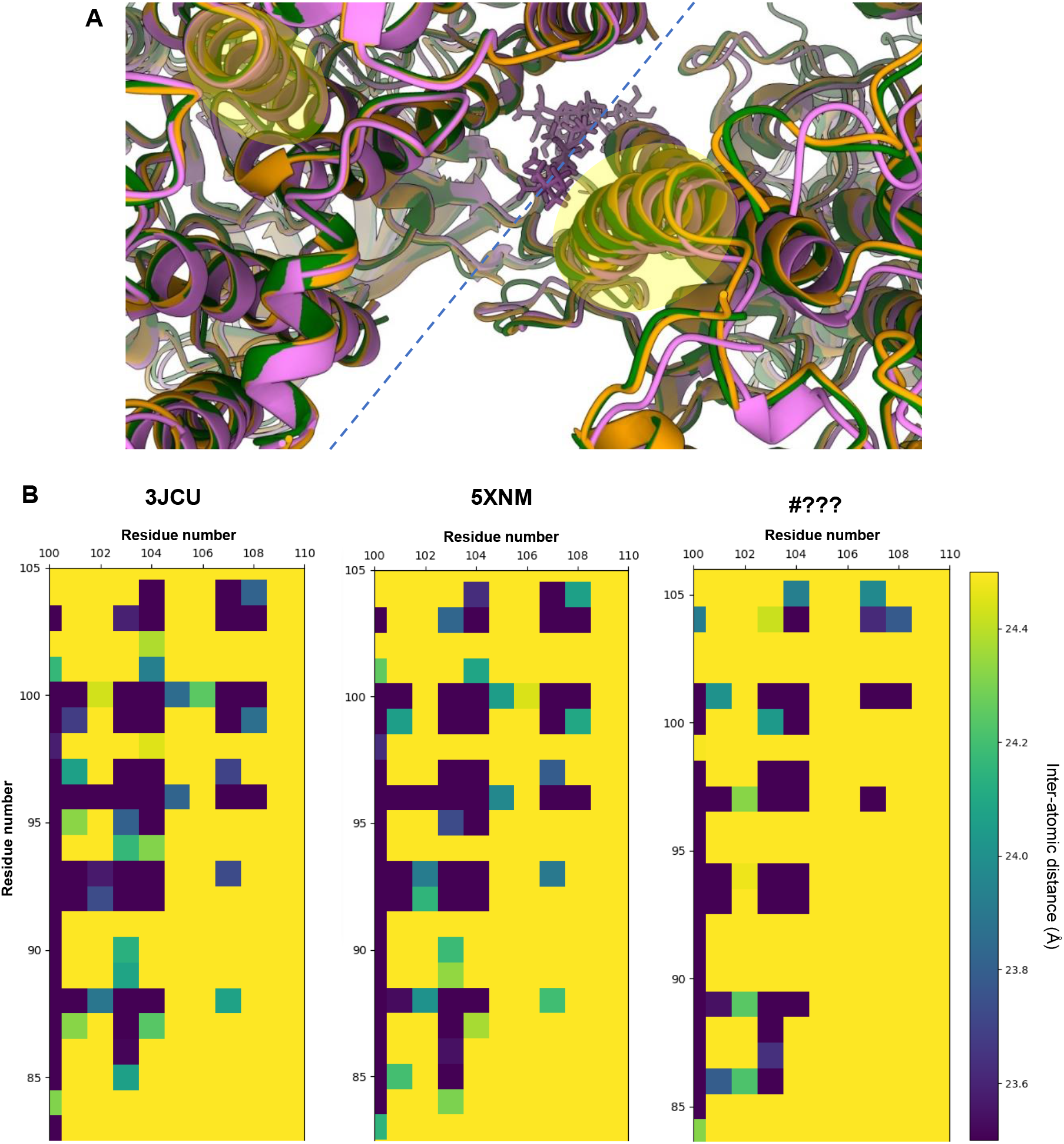
**A**, Dimerization interface between monomeric PSII cores: 3JCU in green, 5XNM in orange, #??? in violet. At the dimerization axis (represented in blue dashed line) a digitonin molecule can be found on model #???. Models were aligned to the D1 protein of the same monomer (to the left of the dimerization interface) **B**, Heatmaps reporting the calculated intermonomer Cα-Cα atomic distances, for each high-resolution higher plant PSII model, between α-helix B of the D1 protein (xx axis) and α-helix B of the CP47 protein from the opposite PSII monomer (yy axis)—highlighted in **A**.

Digitonin molecules have a hexacyclic structure—a lipophilic steroidal sapogenin^48^—that grants partial rigidity to the molecule and result in a high-level of occupancy in the EM map (Appendix, Fig S7). In contrast, the hydrophilic oligosaccharide chains of digitonin stick outside the membrane plane leading to a higher degree of flexibility becoming practically unresolved.

The broadly accessible luminal pocket at the centre of each S-LHCII trimer contains at least seven digitonin molecules (Appendix, Fig S12) that destabilise the natural configuration of the LHCII complexes. We found that the LHCII trimers lack the violaxanthin pigments—ligand code XAT1622—reported to be located at the interface between the monomers^20,22^. In each Lhcb monomer, a digitonin molecule was found to disrupt the interaction between the violaxanthin terminal ring located on the inner side of the LHCII trimer and the chlorophyll *b* CHL607. The opposite terminal ring of the violaxanthin molecule—facing the outside-side of the LHCII trimer—is known to interact with the phosphatidylglycerol (PG) lipid LHG2630. Studies show that LHG2630 plays an essential role in the trimerization of Lhcb proteins^49^. Our EM map indicates that the hydrophobic chains of LHG2630 adopt a new conformation while the molecule remains tightly bound to the complex (Appendix, Fig S12). This interaction is sustained by hydrogen bonds and an ionic interaction between the PG head group and the amino acids Tyr78 and Lys217, respectively. Our structural data confirms that this phospholipid interacts with chlorophyll *a* molecule CLA611 through coordination between its phosphodiester group and the chlorophyll central magnesium atom. Studies of PSII dimerization show that the trans-hexadecanoic fatty acid chains of PG and not its phosphodiester group are the essential factor at the oligomerisation interface^50^. Our finding suggests that one should not infer the same conclusions for PG lipids at other positions of the

PSII complex. The head group of LHG2630 lipids is crucial to stabilise the monomer-monomer interfaces in LHCII and ligates CLA611. CHL601, which in Pea and Spinach PSII structures is also in contact with XAT1622, remains bound to PSII.

Further investigation of our EM map allowed us to observe that a digitonin molecule occupies the binding site of the digalactosyldiacylglycerol (DGDG) lipid DGD520 found in Pea and Spinach PSII (**Fig 2B**). Due to the loss of interaction with DGD520, the neighbouring galactolipid DGD519 has a high degree of flexibility. In Pea PSII, DGD519 hydrogen bonds with DGD520 and the CP43 loop E (Leu401-Ala416), which explains the flexibility of this loop (**Fig 5**). Furthermore, the digitonin-replaced lipid, DGD520, is responsible to mediate the connection between the PsbJ C-terminus and the PSII core through hydrogen bond interactions. This structural finding can explain the loss of PsbJ subunit in our structure.

**Figure 5.**
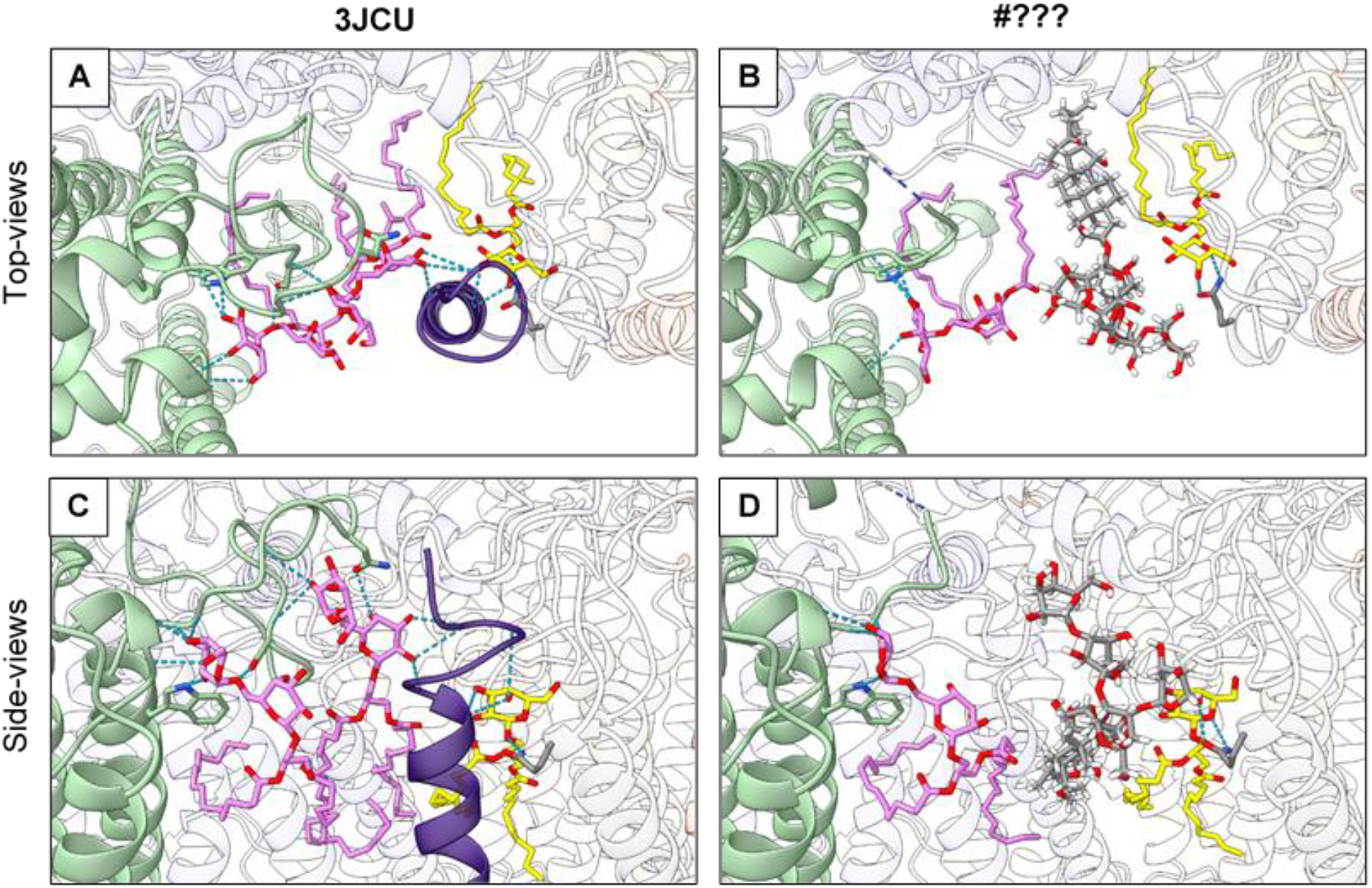
Hydrogen-bonding between loop E of CP43 (in light green) and PsbJ (in purple) subunit are mediated by MGDG (in yellow) and PG (not represented) lipids and disrupted by a digitonin molecule. A and B are top-views, while C and D represent views along the membrane plane, centered around the DGD520 lipid (in pink) present in 3JCU, but replaced by a digitonin molecule in #???. Hydrogen bonds are represented by dashed blue lines and the interrupted loop E of CP43 is represented by a dashed-line half-coloured in blue (N-terminus) and light green (C-terminus).

It should be noted that the cofactors LHG2630, XAT1622, DGD519 and DGD520 were not observed in the 5MDX model.

### The PsbO short-loop is disordered as result of the digitonin bound between PSII monomers

The extrinsic protein PsbO has two loops—here termed short-loop (Ala154 to Pro156) and long-loop (Ser239 to Ala279)—to which structural information have been difficult to obtain for Arabidopsis PSII (Appendix, Fig S13). In Spinach PSII^22^, which retains the extrinsic PsbP and PsbQ proteins, both loop regions are well structured. The unstacked PSII structure from Pea (PDB: 5XNM) has the short-loop modelled but the long-loop, which is contiguous to PsbP in Spinach PSII and in Pea stacked PSII (PDB: 5XNL) models, is absent.

The increased quality of our EM map revealed three additional amino acids in the short-loop N-terminal with a conformation dissimilar to their counterpart amino acids in Pea and Spinach models. The amino acid sequence of the short-loop peptide is the poorest conserved region of the PsbO protein among higher plants (Appendix, Fig S14), suggesting that this flexible region is of lower functional relevance. Nevertheless, it is conserved to a higher degree in Spinach and Pea, while being the least conserved in Arabidopsis PsbO, justifying the diverged structural arrangement (Asn153-Glu157, Appendix, Fig S15) of the peptide backbone including the newly modelled amino acids. The hydrophilic sugar chains of the digitonin molecules harboured between PSII monomers are within no distance to interact with PsbO short-loop, and thus could be the reason for its disorder.

To note that we do not exclude the possibility that an even larger number of detergent molecules are bound to the complex, even if they cannot be observed due to their high flexibility.

### The disordered PsbO long-loop is related to several structural changes at the OEC

In the here presented PSII structure, the C-terminus of the D2 protein was found to be twisted (Appendix, Fig S16) in a similar manner to what was reported for the cyanobacterial *Synechocystis* Apo-PSII^40^—making the last C-terminal residues occupy the region where the otherwise ordered PsbO long-loop would be located.

The PsbO-Asp243 (Asp158, according to 5XNM sequence numbering) is a residue of PsbO long-loop known to participate in a hydrogen-bonding network that connects the Mn_4_CaO_5_ to the lumen and stabilise the binding of PsbP and PsbQ proteins^51^. Interestingly, the 5XNM model presents the long-loop modelled to a higher extent (six additional amino acids at the C-terminus, including Asp158). Our analyses of hydrogen-bonding protein-protein interactions of the 5XNM model show that Asp158 and Leu157 hydrogen-bond residues D1-Arg334 and D1-Asn335, respectively. These interactions are responsible to maintain the structure of the D1 C-terminal and consequently keep the Mn_4_CaO_5_ in its correct position.

Considering that the long-loop region of PsbO is highly conserved in almost all higher plants, we can assume that the observed disorder in Arabidopsis PSII models is linked to the loss of the extrinsic proteins PsbP and PsbQ. Nevertheless, it is unclear why the C-terminus of the D2 protein in our PSII structure is twisted and why the unstacked PSII from Pea has a more ordered PsbO long-loop, and a structured D1 C-terminus capable of binding the inorganic cluster. We hypothesize that both observations are correlated to each other, and a possible consequence of the use of digitonin as a solubilization agent.

### The OEC in the absence of PsbP and PsbQ proteins

The Mn_4_CaO_5_ cluster is coordinated by crucial amino acids of the D1 protein and the core antennae protein CP43, often referred to as “first shell” residues. Amino acids from the D2 protein and the inner antenna CP47 are essential “second shell” residues that: sustain the structure of the OEC and form the channels which deliver H_2_O molecules to the water oxidation cluster. The extrinsic subunits are known to stabilise the hydrophilic pocket that hosts the inorganic cluster. Comparing our model to the 3JCU mature PSII structure containing the extrinsic proteins and the Mn cluster, we found that in the absence of PsbP and PsbQ proteins, the Mn_4_CaO_5_ binding pocket undergoes conformational changes. Without the presence of the Mn_4_CaO_5_ cluster, the C-terminal tail of the D1 protein (amino acids His337-Ala344) was found to be disordered and therefore not visible in our EM map. Out of the seven amino acid that ligate to Mn_4_CaO_5_, only two, D1-Asp170 and D1-Glu189, retain their relative positions to the mature Spinach PSII structure (**Fig 6A**). D1-Glu329 adopts an unusual conformation, where it is found to hydrogen bond with D1-His332. Glutamate 333 of the D1 protein, one of the coordinating residues of the manganese cluster, is consequently in a different conformation (**Fig 6B**) and hydrogen-bonded to D1-Glu329. The C-terminus of the D1 protein (amino acids 337-344) is disordered and not visible in our EM map.

**Figure 6.**
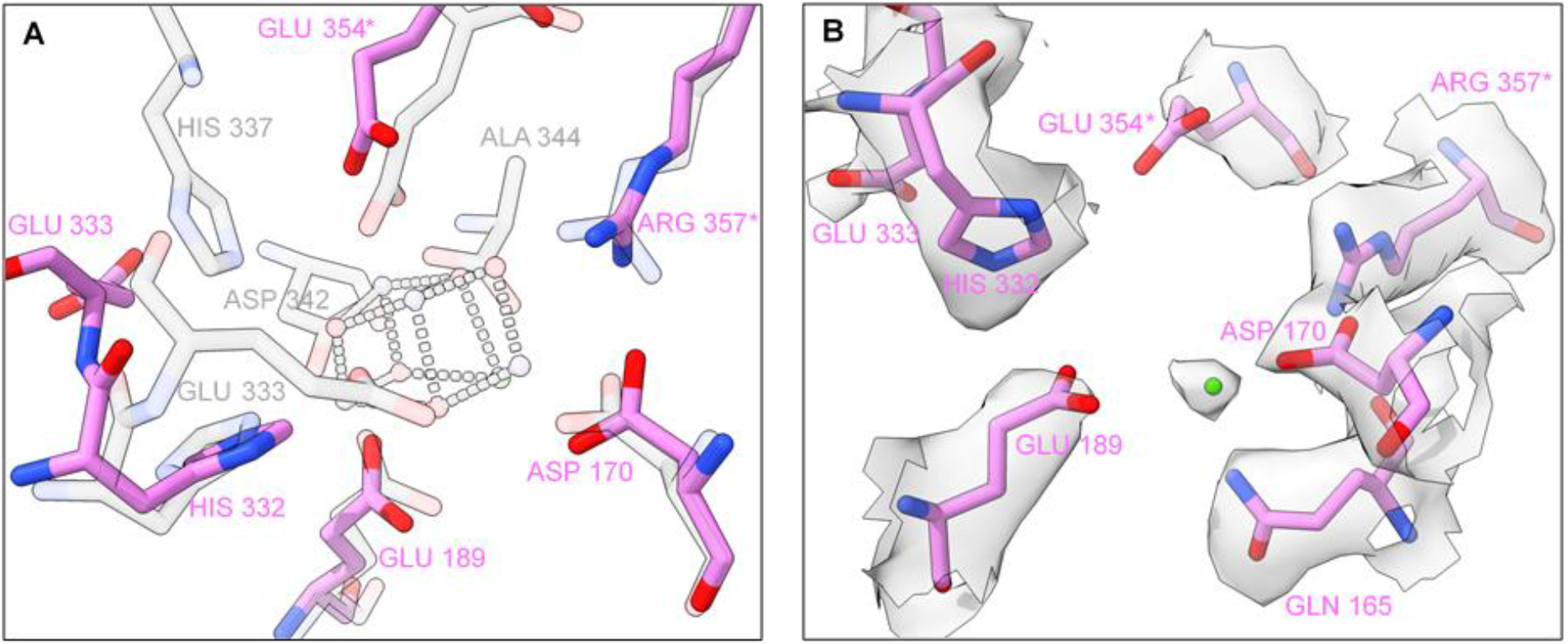
The manganese cluster binding site. A, Amino acid ligands to Mn4CaO5: Spinach PSII (3JCU, transparent grey) superimposed to Arabidopsis PSII (#???, pink). To avoid double labelling, the overlapping residues of 3JCU are not identified for clarity. B, Arabidopsis amino acid ligands to Mn4CaO5 fit into their respective densities and an extra density attributed to a positive metal ion (in our structure modelled as manganese). *Identifies the amino acids belonging to the CP43 protein, while the remaining apply to the D1 subunit.

The core antenna protein CP43 consists of six transmembrane helices connected by short loops (loop A-D) and a very large extrinsic domain known as loop E. Glutamate 354 and arginine 357, also ligands of the Mn_4_CaO_5_, are part of loop E. Both amino acids nearly conserve their position relative to mature PSII (**Fig 6A**). The PsbO seems to play an essential role in stabilising the CP43 loop E—connected by 10 interprotein hydrogen bonds—that otherwise would have high degree of flexibility. Finally, near the water oxidation cluster binding site, loop E is stabilised via hydrogen bonds between CP43-Arg357 and D1-Asp170. A careful analysis reveals that CP43-Glu354 is slightly receded from its coordinating position to Mn2 and Mn3 of the Mn_4_CaO_5_, but pre-organised to receive and coordinate the manganese ions (**Fig 6A**). At the expected position of Ca^2+^ ion on a mature PSII structure, we find a density that can be attributed to a positively charged ion (**Fig 6B**), potentially coordinated by the D1 residues Gln165 (also retaining its position related to mature PSII), Asp170 and Glu189. EM data by itself offers no signifying conclusion regarding the identity of the metal ion, which could be a Ca^2+^ ion, or the first manganese ion that binds to PSII to form the Mn_4_CaO_5_. If the latter is true, this binding site might well be the long sought high-affinity site of PSII oxygen-evolving complex.

## Discussion

In this work, we presented the first high-resolution structure of Photosystem II from wild-type *Arabidopsis thaliana*, allowing a detailed comparison to other higher plants’ PSII structures, while fairer comparison to existing high-resolution cyanobacterial PSII models. The quality of the obtained EM map enabled a better and more comprehensive modelling of most of the PSII core proteins; a detailed analysis of the positioning of chlorophyll, carotenoid, and lipid molecules; and the placement of water molecules, of which some compose the water channels delivering substrate to the catalytic site of this water-splitting enzyme.

To isolate PSII, we solubilised thylakoid membranes with a detergent mixture containing β-DDM—a harsher isoform of the more commonly used α-DDM^32^—and digitonin. Even though β-DDM can be a more disruptive detergent hindering the obtention of large PSII supercomplexes, we did not find evidence of any structural changes caused by it. Instead, we found that digitonin molecules disrupt several regions of PSII by replacing lipids that keep the structural integrity of the membrane protein complex. Exploiting the digitonin effects on the major antennae, we found that the PG phosphodiester group is the essential lipid element to stabilise the trimerization of LHCII complexes. Furthermore, we found that digitonin molecules are responsible for: the loss of the PsbJ subunit; the consequent instability of a CP43 loop; the disordered D1 C-terminus and the resultant fall of the Mn_4_CaO_5_ cluster. Besides revealing new knowledge, our model is a representation of the disruptive mechanisms that digitonin molecules can have in the stability of protein complexes, leading to a dramatic loss of function.

The PsbJ protein is one of the most hydrophobic proteins in the thylakoid membrane and has been suggested to be involved in the electron flow within PSII^52^ even though it does not seem to bind any pigment or cofactor. It is found in Pea and Spinach structures^20,22^ bound to the PSII complex through hydrogen bonding with lipids associated with CP43 protein, but neither found in our Arabidopsis PSII structure, nor in the 5MDX model. Regarding the PsbY protein, we do not detect any densities suggesting the presence of this low molecular mass subunit— as for the PSII models of Spinach (3JCU) and Pea (5XNM and 5XNL)—even when western-blotting analysis show that the protein is present in various PSII samples from higher plants including Arabidopsis^53^. So far, the structure of the PsbY subunit has only been observed on cyanobacterial PSII models^29^, which in higher plants C_2_S_2_M_2_-type PSII would be located on the outskirts of the complex (considering a conserved position). The small proteins PsbY and PsbJ are both hydrophobic, have a single transmembrane helix and are suggested to exist in the outskirts of the complex, thus likely to be dissociated from the complex in the presence of detergents.

As previously suggested^54^, the low molecular mass protein PsbJ might be required to stably bind PsbR, which in turn is responsible for protecting PsbP and PsbQ from proteolytic degradation. PsbO is known to be the most stable extrinsic protein, weakly sensitive to the depletion of the neighbouring PsbP and PsbQ proteins. We anticipate that the PsbO long-loop requires PsbP to maintain its stability, which is correlated with the loss of the transmembrane subunits PsbJ and PsbR, and the extrinsic proteins PsbP and PsbQ. Furthermore, the oxygen-evolving proteins PsbP and PsbQ are pH-sensitive^20^, and both proteins bind the strongest to the PSII lumen side around pH 6^31^. Unfortunately, PSII complexes tend to aggregate at pH 6, obstructing single-particle cryo-EM studies. The isolation of PSII complexes at pH 6 using alternatives to detergents, such as amphiphilic^17^ or styrene-maleic acid (SMA) copolymers forming membrane proteins embedded in nanodiscs, might offer a solution to these challenges. Not less crucial, such copolymers would protect PSII from the undesirable effects of detergents and allow to extract PsbJ (and possibly PsbR and PsbY) bound to PSII.

In comparison to other PSII structures, new knowledge on the assembly of the OEC from higher plants can possibly be derived from our model which exhibits similar characteristics to PSII still in assembly stage, prior to the binding of the extrinsic PsbP and PsbQ: the monomeric *Synechocystis* apo-PSII^40^ presents analogous features to the high-resolution Arabidopsis PSII, particularly at the Mn_4_CaO_5_ binding-site.

These findings allow us to step forward in PSII research and bring new research questions: Could the detergent molecules anchored inside the moiety of isolated chlorophyll-binding proteins alter their spectroscopic properties; how has that influenced the outcomes of different PSII and LHCII spectroscopical studies; and if digitonin molecules fit into the centre of the LHCII trimer, did they replace other naturally present molecules such as yet undetermined lipids at those positions?

## Materials and Methods

### Photosystem II supercomplex extraction

*Arabidopsis thaliana* (ecotype Columbia-0) plants were grown on soil in greenhouse at Umeå University (Umeå, Sweden) for 8 weeks under controlled temperature (23°C) and humidity (70%) conditions, with 8 h light / 16 h dark photoperiod. Seeds were obtained and identified by Arabidopsis Biological Resource Center (ABRC; https://abrc.osu.edu/). Local and national regulations was obeyed and the permission from Jordbruksverket - Växtkontrollenheten has been obtained (Dnr 4.6.20-6365/15).

Plants were harvested after a dark period of 16 h and BBY membranes^41^ prepared according to previous protocol^55^. The extraction of PSII C_2_S_2_M_2_ complexes was performed^31^ with the following modifications: solubilisation was performed by adding a detergent mixture of 0.2% β-DDM and 1% digitonin, followed by a 30 min dark incubation at 4°C.

### Sample preparation and electron microscopy

For cryo-electron microscopy, solubilised PSII sample was washed with a 10mM HEPES-KOH pH 7.5 + 0.01% w/v digitonin solution (bellow CMC level) and concentrated to 1 mg/ml Chl using an Amicon Ultra 30k centrifugation filter (Millipore). 4 μl concentrated sample was applied to glow-discharged (30 s at 50 mA) Quantifoil R 1.2/1.3 Cu 300 grids (Quantifoil Micro Tools) and grids were plunge frozen in liquid ethane using an FEI Vitrobot MkIV (Thermo Fisher Scientific) at 100% humidity, 4°C, using a blot force of −5, wait time of 1 second, and blotting time of 5 seconds. Automated data collection was performed using EPU software on a Titan Krios G2 transmission electron microscope (Thermo Fisher Scientific) operated at 300 kV, equipped with a Gatan K2 Summit direct electron detector and Gatan Quantum GIF energy filter at the Umeå Core Facility for Electron Microscopy, a node of the Swedish National Cryo-EM facility. In total 12227 movies, fractionated in 40 frames, were collected at a pixel size of 0.82 Å, total dose of 59.7 e^−^/Å^2^ and defocus range of -1.5 to -3.0 μm. On the same microscope and grid, a data set was collected using Volta phase plate^56^: in total 3585 movies, fractionated in 40 frames, were collected at a pixel size of 1.09 Å, total dose of 36.7 e^−^/Å^2^ and defocus of -0.7 μm.

### Cryo-EM data processing, model building and refinement

The collected datasets were processed using the software package cryoSPARC^57^ (version v2.15).

#### VPP dataset

Automated particle picking based on 116 manually picked templates was carried out before the major 2D classification round. The combination of well-resolved 2D classes isolated a subset of 80456 particles, out of 273202 particles automatically picked. The *ab-Initio* 3D reconstruction strategy segregated particles into 2 different 3D-classes, allowing us to proceed with one class of a more homogeneous set of 56322 particles. Homogeneous refinement of the preliminary 3D volume was carried with imposed C2 symmetry, yielding a 3D reconstruction with an estimated average resolution of 3.6 Å.

High-resolution data set: Template-free automated particle picking was carried out before the major 2D classification round. From the selected 2D classes we restricted the data set to a total of 110 659 particles, out of 416 262 particles automatically picked. The *ab-Initio* 3D reconstruction strategy segregated particles into 4 different 3D-classes, allowing us to proceed with a more homogeneous subset of 100 712 particles. The selected particles were individually motion-corrected, and homogeneous refinement of the preliminary 3D volume was carried with imposed C2 symmetry. After a round of per-particle CTF refinement, the final homogeneous refinement yielded a 3D reconstruction with an estimated average resolution of 2.79Å, using the gold standard FSC (Appendix, Fig S1).

Three independent post-processing methods were applied to the resultant high-resolution electrostatic potential map: local sharping with Phenix Autosharpen tool^58^ (Phenix v1.18.2); density modification with Phenix denmod tool^59^ (Phenix v1.18.2) using the outputted half-maps by cryoSPARC; application of the negative of the Laplacian operator using Chimera^60^ (version 1.14). The three resulting post-processed maps were useful to confirm certain densities, such as the presence of ions and water molecules.

For model building, the previous Arabidopsis atomic model 5MDX was manually docked and each chain morph-fitted to the 2.79 Å map using COOT^61^ (version 0.92). The new cofactors and the PsbTn subunit where found using the 5XNM Pea PSII model as a template. Overall the presented model was manually refined using COOT and both the high-resolution and the VPP dataset, and automatically refined with Phenix Real-space refinement tool^62^ (Phenix v1.18.2).

### Model analysis and preparation of figures

Molecular graphics and analyses were performed with UCSF ChimeraX^63^ (version 1.1), developed by the Resource for Biocomputing, Visualization, and Informatics at the University of California, San Francisco. Surface electrostatics were calculated using the software packages PDB2PQR (to assign atomic charges and radii to the protein backbone using force field parameters) and APBS (to generate the electrostatic surface)^64^. We recommend ChimeraX for model and surface visualisation.

## Acknowledgements

The authors thank to Dr Johannes Messinger, Dr Thomas Kieselbach and Dr Domenica Farci for reviewing the manuscript. This work was supported by grants from the Carl Trygger foundation to WPS, and the Swedish Research Council (VR), grant 2016-05009 to KP. The data was collected at the Umeå Core Facility for Electron Microscopy, a node of the Cryo-EM Swedish National Facility, funded by the Knut and Alice Wallenberg, Family Erling Persson and Kempe Foundations, SciLifeLab, Stockholm University and Umeå University.

## Author contributions

A.T.G. and W.P.S. designed research; A.T.G performed sample preparation, data processing, and model building; A.T.G and M.H. performed cryo-EM data collection; A.T.G and W.P.S. wrote the paper; M.H. and K.P. assisted model building and writing of the paper.

## Additional information

The executed work with plant material complies with the local and national regulations. The authors declare no competing interests.

## Data availability

The raw multi-frame micrographs of the higher-resolution data set and the VPP dataset were deposited in the Electron Microscopy Public Image Archive with accession codes EMPIAR-##### and EMPIAR-#####, respectively. Electrostatic potential maps used for refinement, unsharpened EM map, half-maps and other relevant post processed maps have been deposited in the Electron Microscopy Data Bank, EM-####. The modelled structure is available in the Protein Data Bank (PDB) under the accession code #???.

