## Supplemental Figures for "High-resolution model of Arabidopsis Photosystem II reveals the consequences of digitonin-extraction"

A

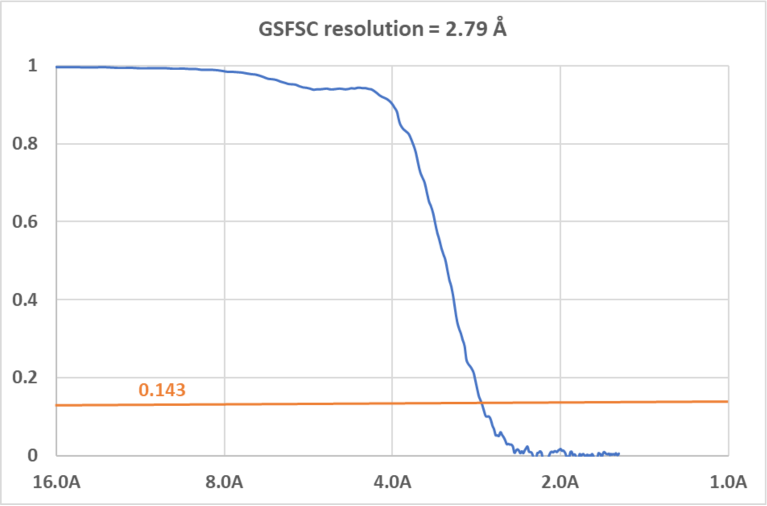


B

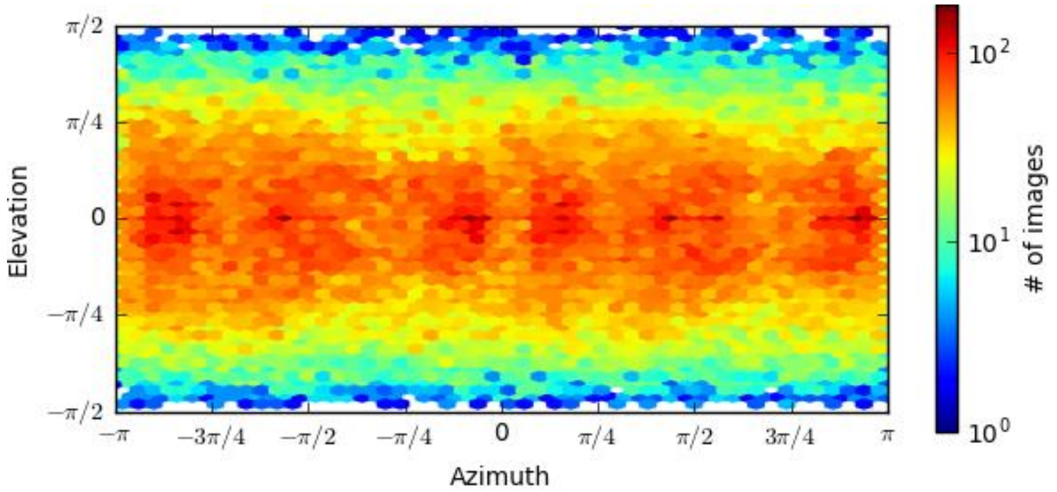


Figure S1 – A, Fourier Shell Correlation (FSC) curve of two independent refined datasets based on the golden-standard criterion for the $C_2S_2$ supercomplex. B, Plot for the angular distribution of particles that participated in the 3D reconstruction of the higher-resolution map.

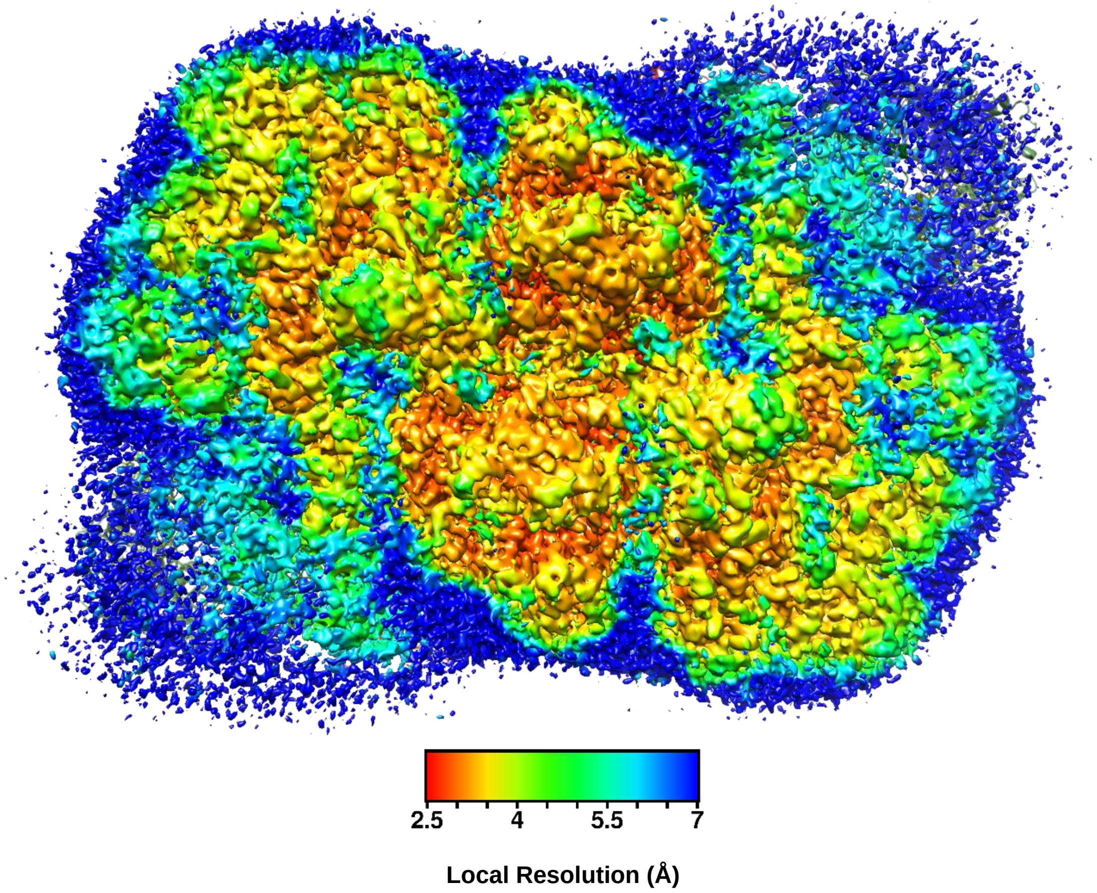


Figure S2 – Top-view (stromal side) of our EM Map coloured according to local resolution. Local resolution was computed using cryoSPARC.

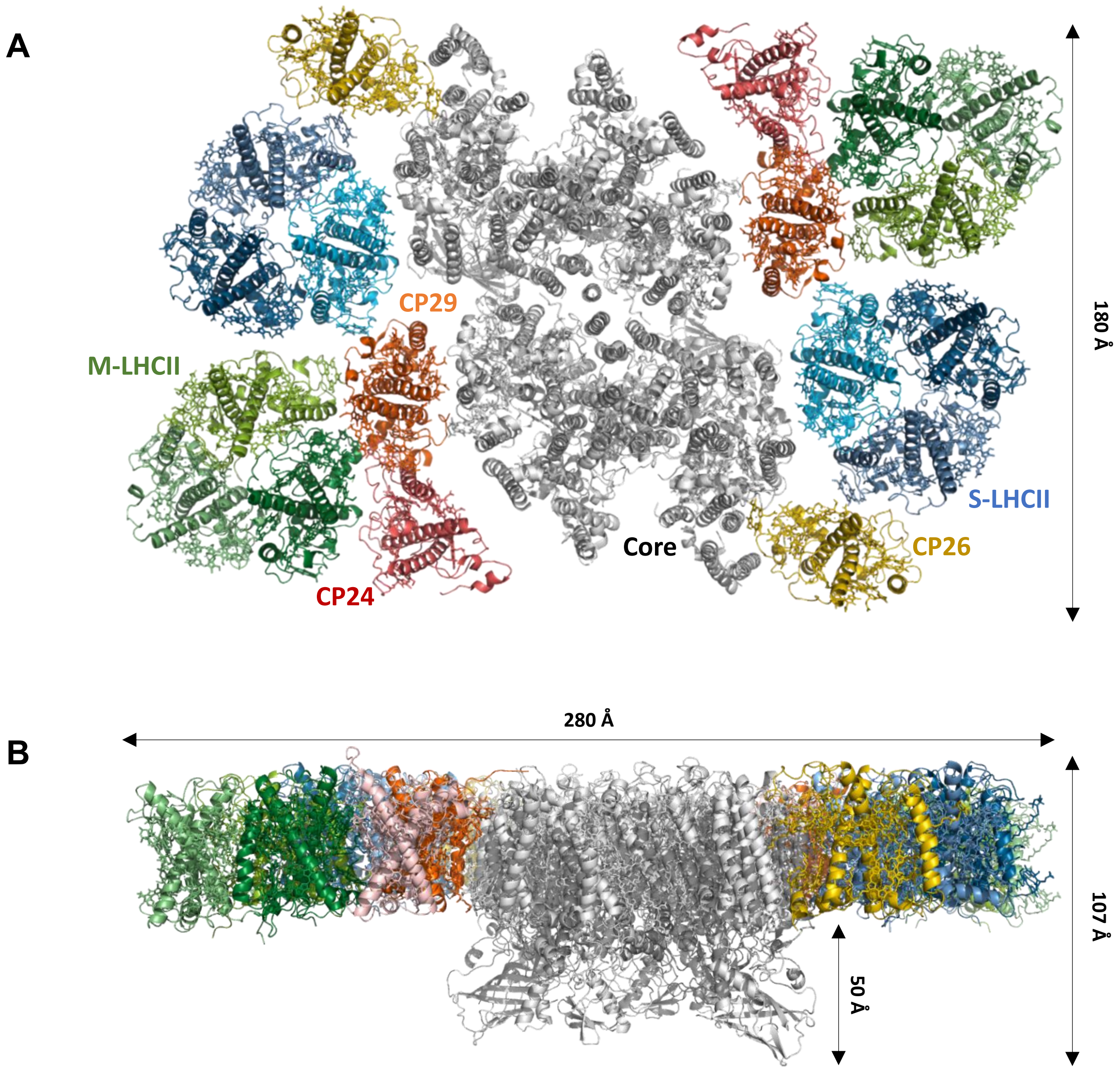


Figure S3 – Overview of the atomic model built from the single-particle Cryo-EM reconstruction of *Arabidopsis thaliana* Photosystem II $C_2S_2M_2$ supercomplex, obtained from electron density map with overall resolution of 3.13 Å: A, top-view of the complex from the stromal side; B, side-view of the complex with the lumen side pointing down.

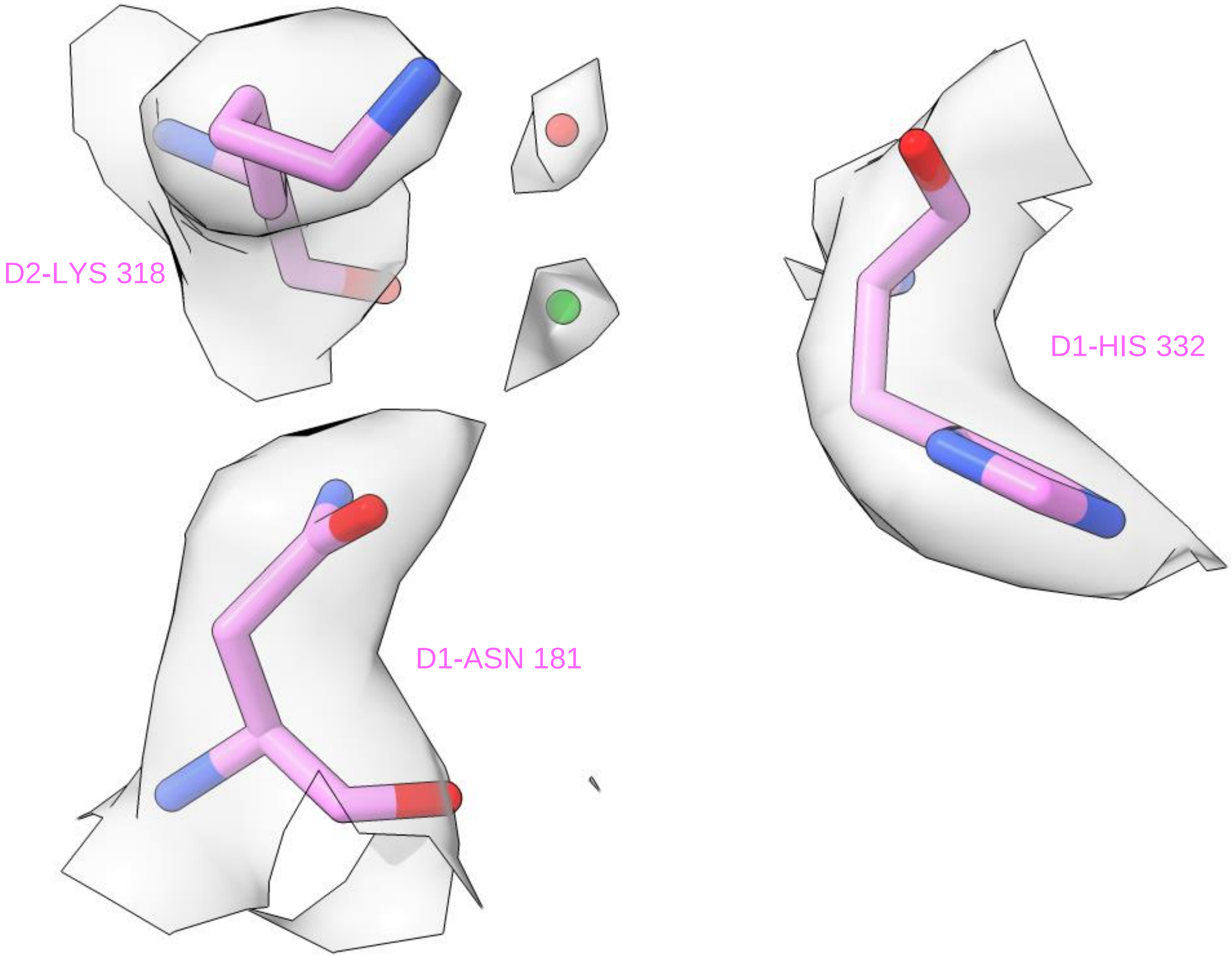


Figure S4 - The binding site of the chloride anion (green) found in our EM map (surface coloured in grey).

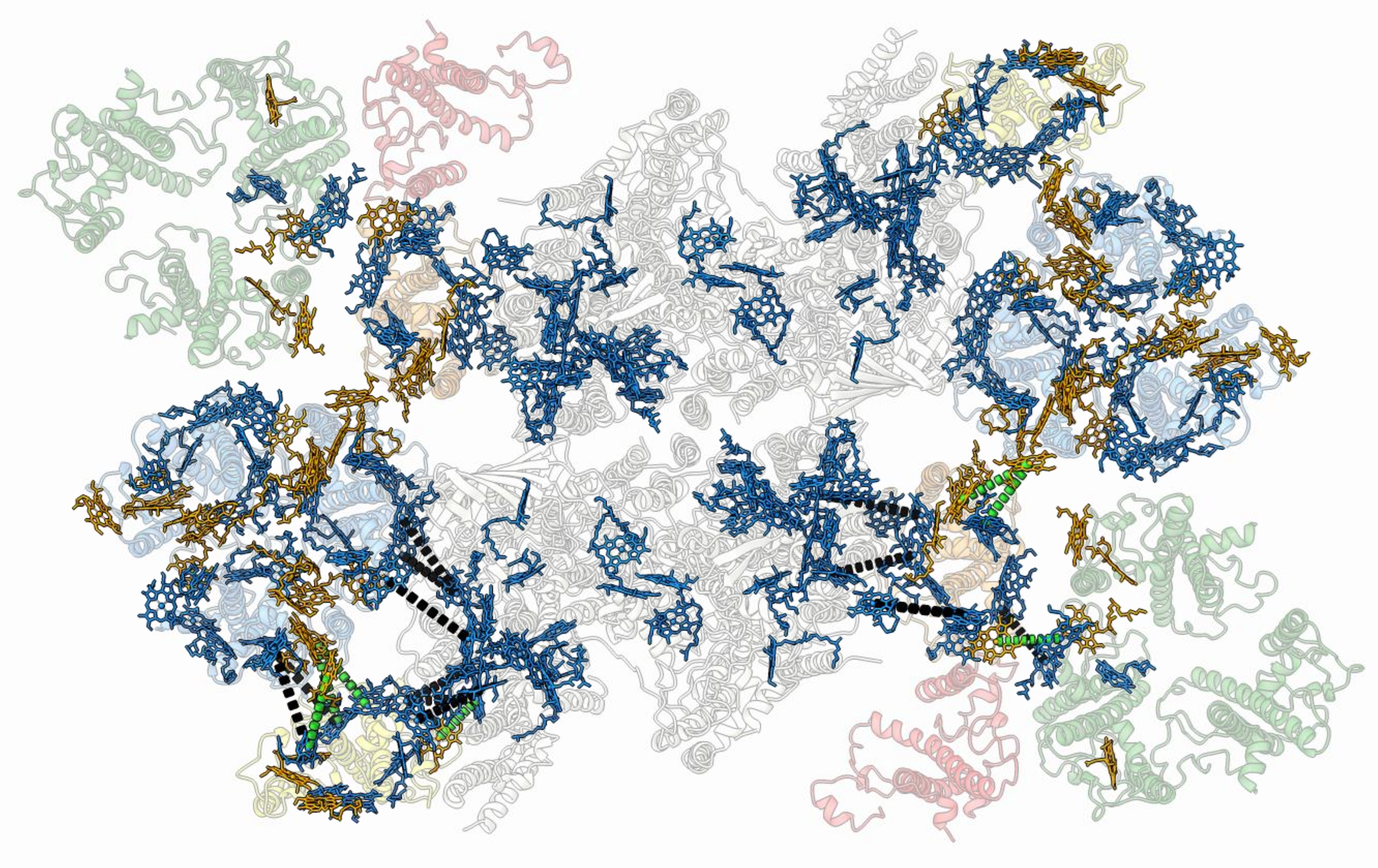

Figure S5 – Top-view of the distribution of chlorophylls in Arabidopsis PSII: chlorophyll *a* are coloured in blue and chlorophyll *b* in orange; the dashed lines represent the energy transfer between two couplings of chlorophyll *a* (black) and a coupling containing at least one chlorophyll *b* (green), corresponding distances can be seen in Table S6.

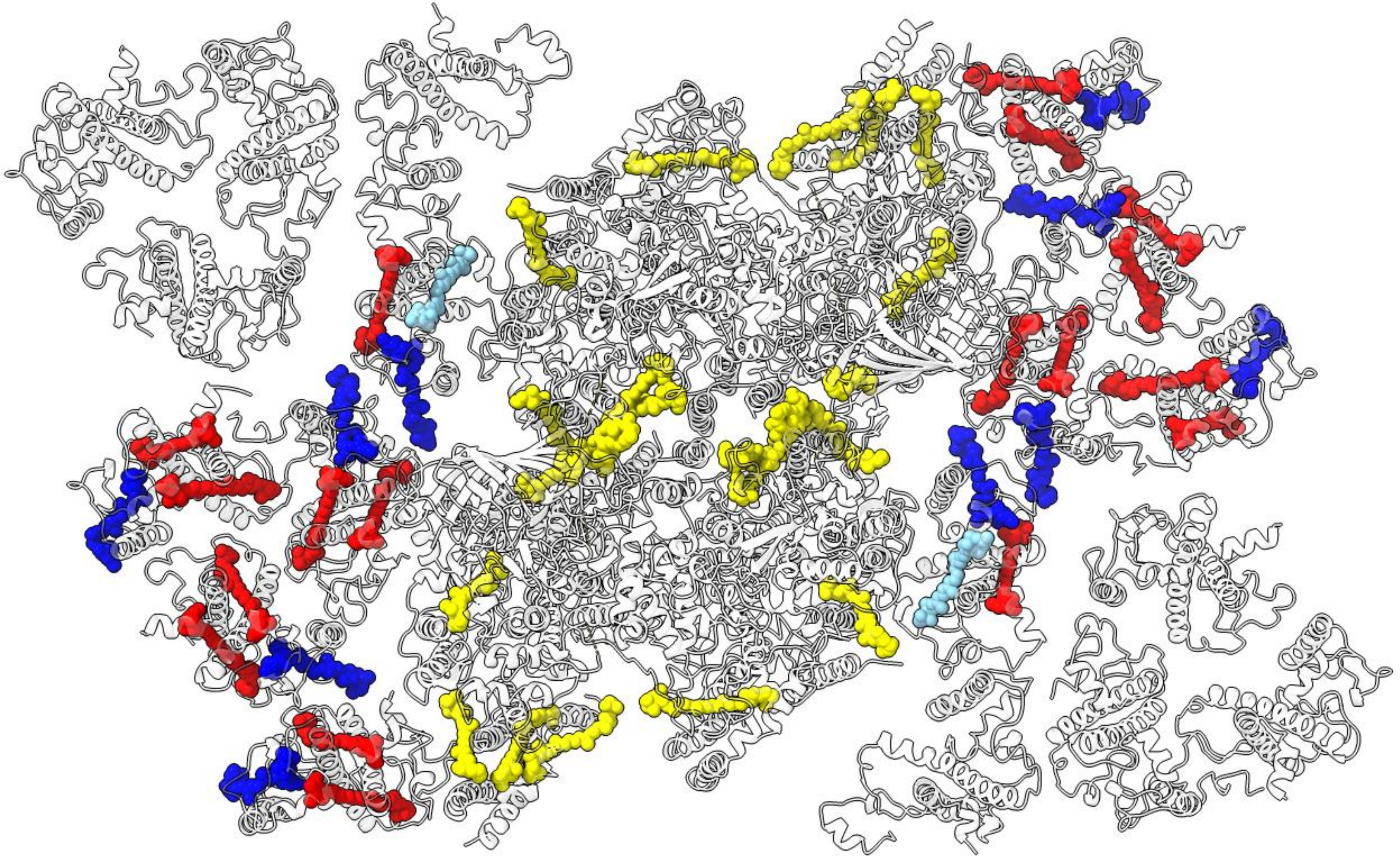

Figure S6 – Top-view of the distribution of carotenoid molecules in Arabidopsis PSII: beta-carotene (yellow), lutein (red), violaxhantin (light blue), neoxantin (blue).

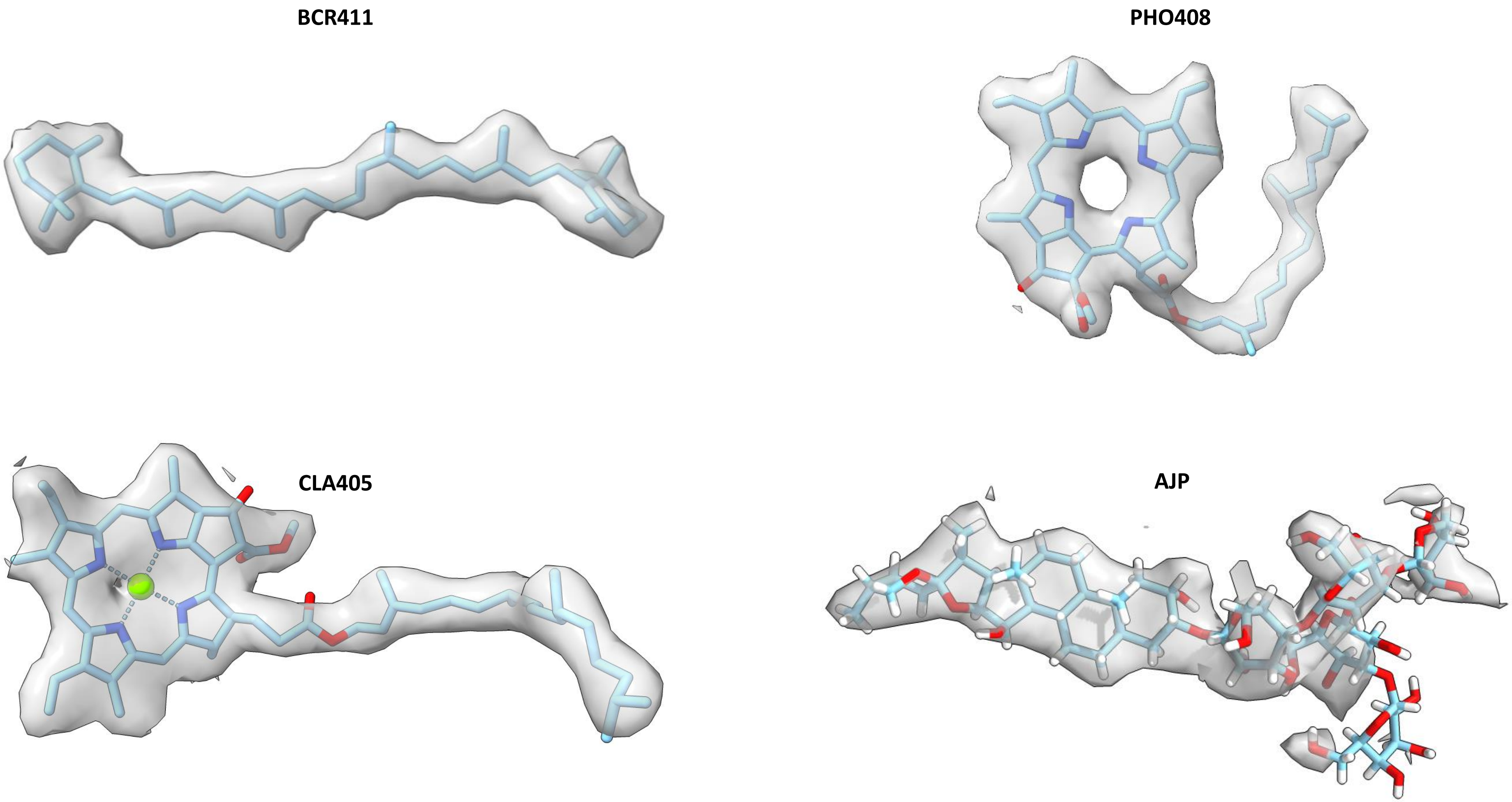


Figure S7 – Exemplars of some modelled ligands fitted into their respective densities: beta-carotene (BCR411), chlorophyll a (CLA405), pheophytin (PHO408) and a digitonin (AJP) molecule.

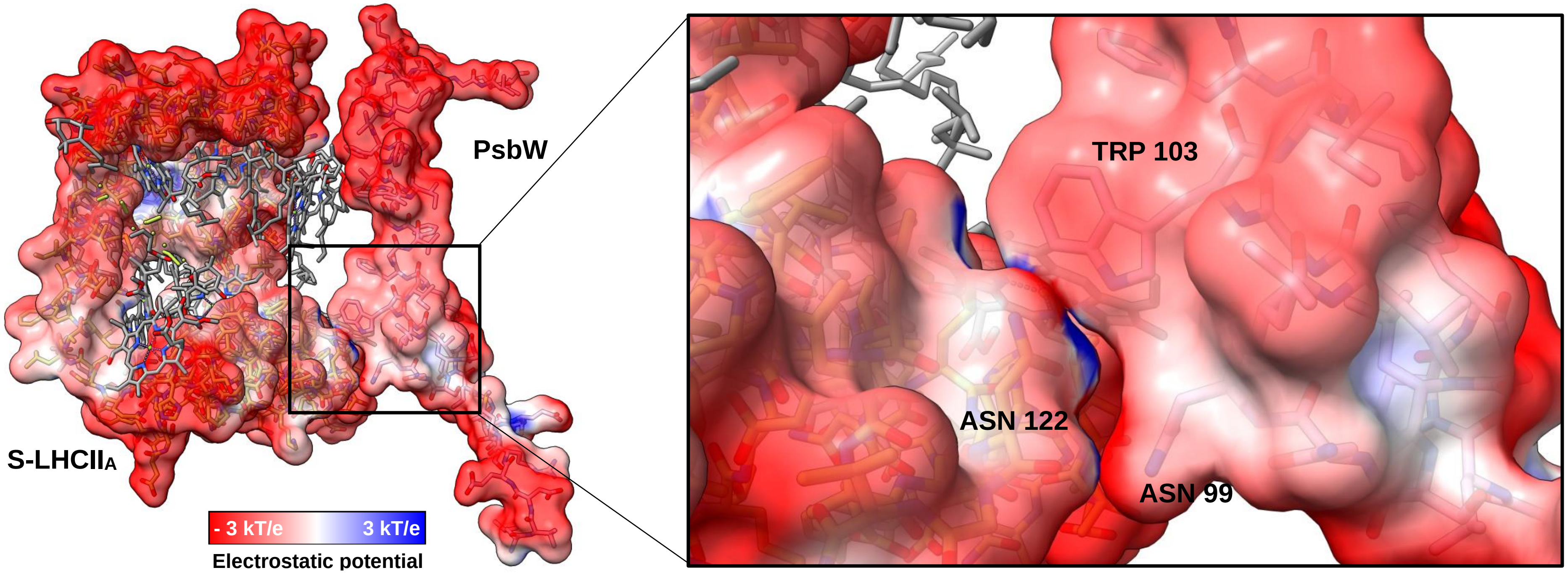


Figure S8 - Electrostatic interactions between PsbW and monomer A of S-LHCII. The atomic model of PsbW and monomer A of S-LHCII, as well as the corresponding surface electrostatics of the polypeptide chains (slightly transparent to aid interpretation) are displayed.

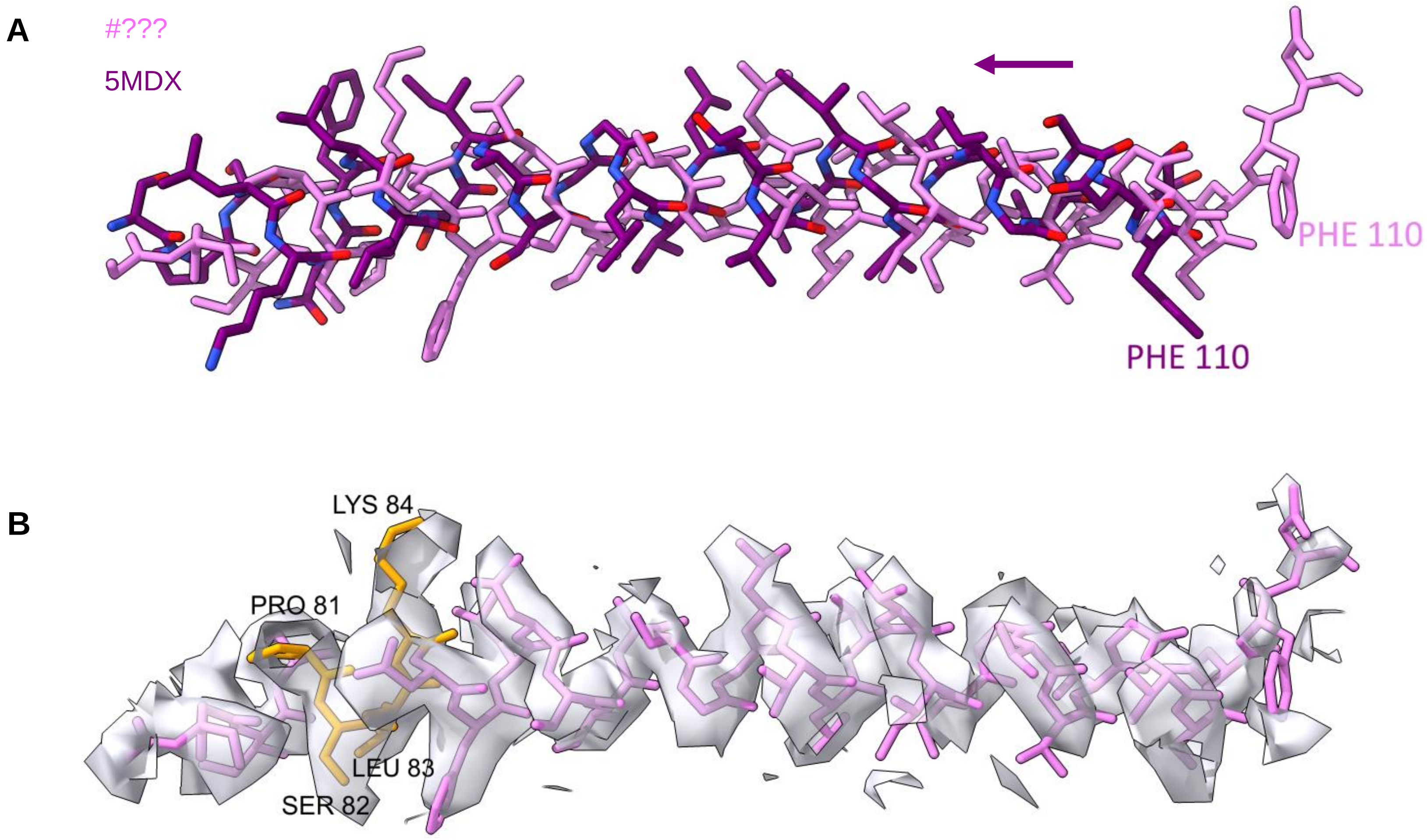


Figure S9 – PsbX: A, representation of the shift between PsbX subunit in our EM model (#???, pink) and the 5MDX model (purple); B, the PsbX subunit fit into its respective density of our EM map.

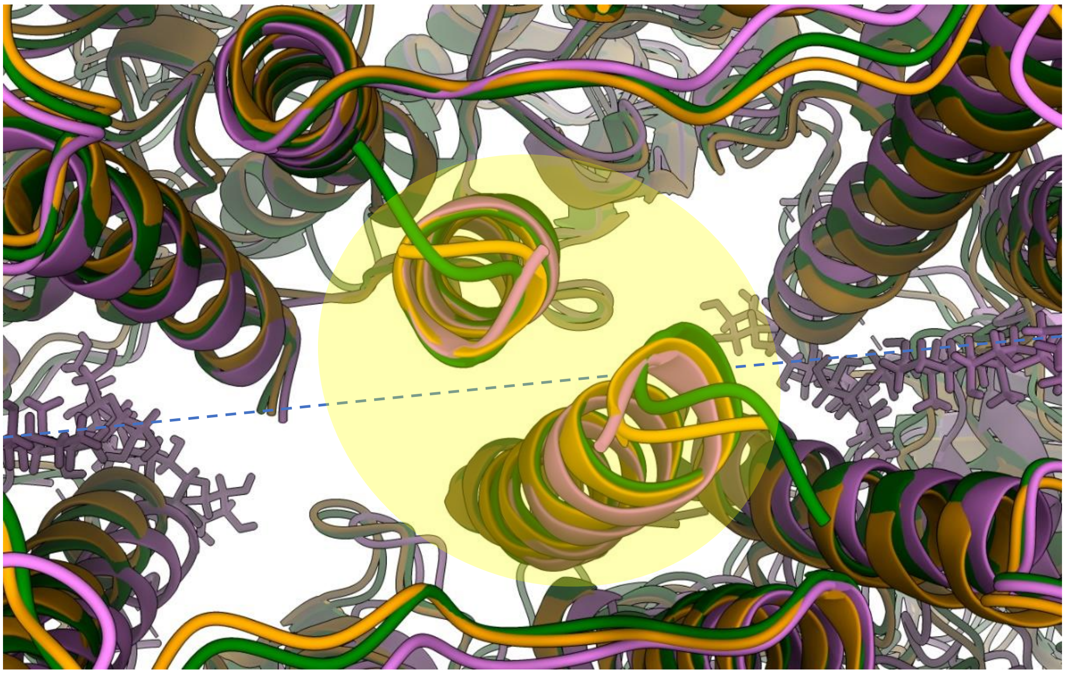

Figure S10 - Dimerization interface between monomeric PSII cores: 3JCU in green, 5XNM in orange, #??? in violet. At the dimerization axis (represented in blue dashed line), two digitonin molecules can be found on model #???. Models aligned to the PsbM protein of the same monomer (above the dimerization interface). The transmembrane helices highlighted in yellow indicate the secondary structure motifs to which intermonomer Cα-Cα atomic distances where calculated (see Figure S11) and the blue dashed lines represent the dimerization axis of the PSII complex.

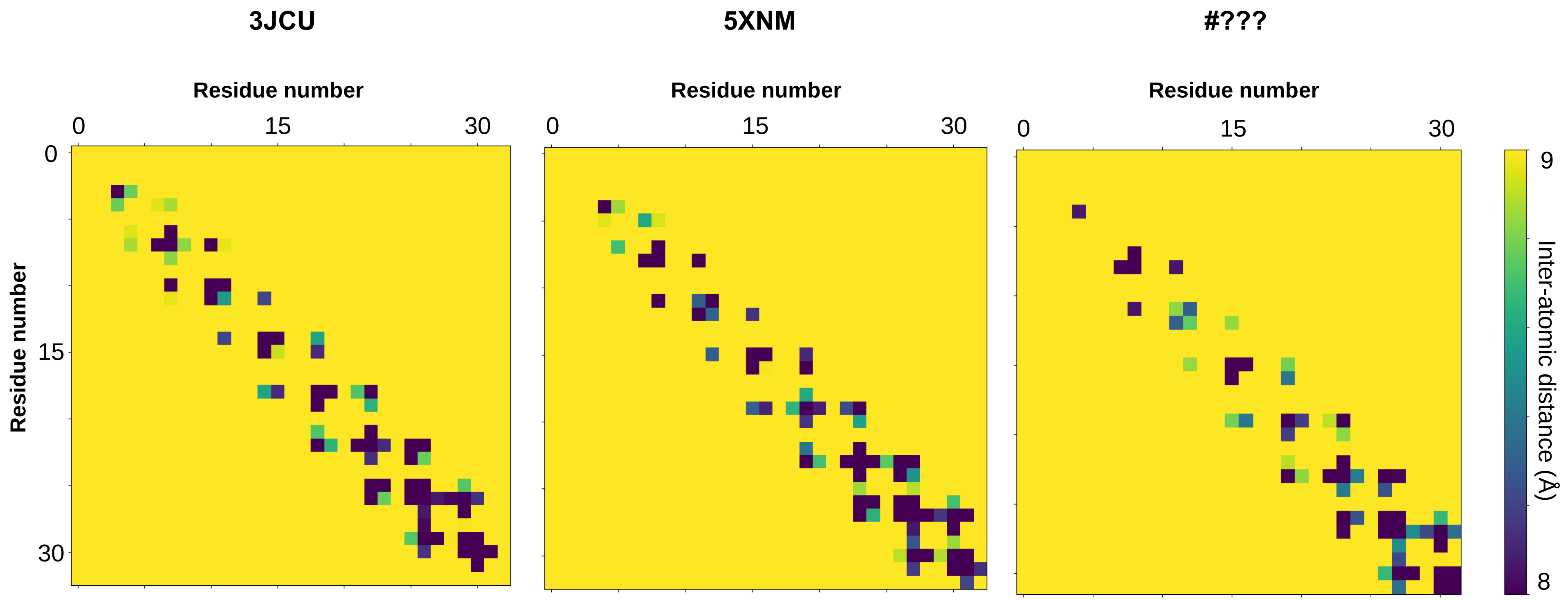


Figure S11 - Heatmaps reporting the calculated intermonomer Cα-Cα atomic distances, for each high-resolution higher plant PSII model, between the small PsbM proteins from opposite PSII monomers—highlighted in yellow in figure S10.

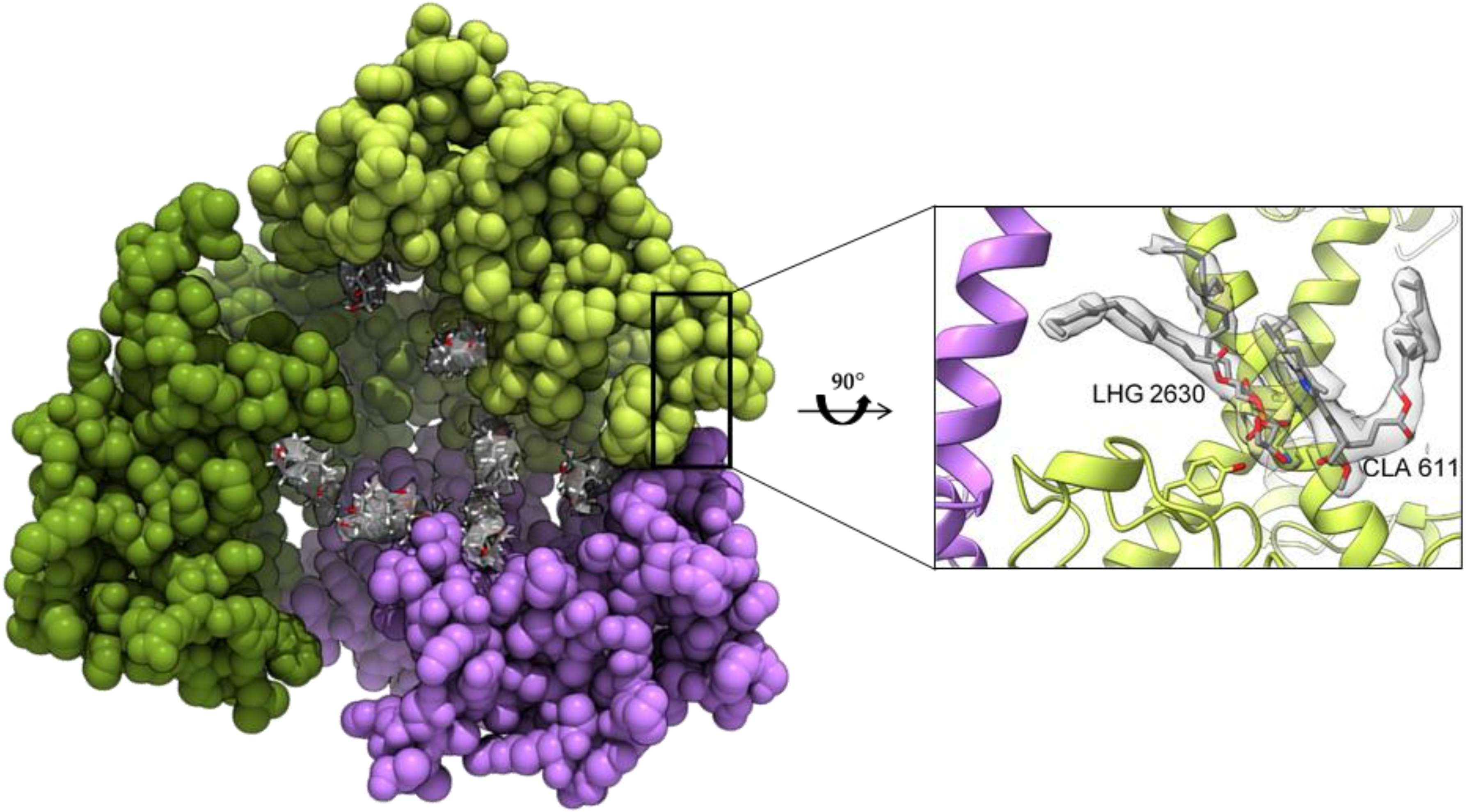


Figure S12 – Digitonin molecules occupy the center of the LHCII trimer. Inset shows the binding site of the PG lipid LHG2630 and chlorophyll a CLA611, bound to the monomer S-LHCIIA. The residues Tyr78 and Lys217 of S-LHCIIA participating in the binding of LHG2630 are displayed.

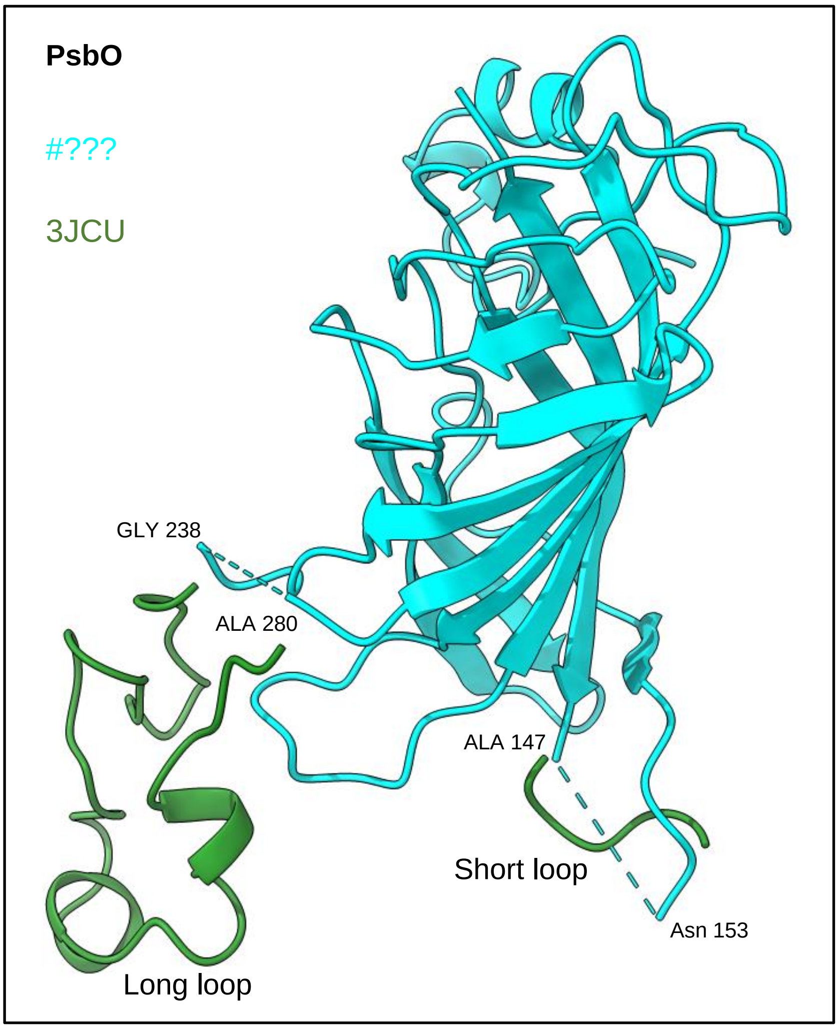


Figure S13 – Representation of PsbO long- and short- loop interruptions (in cyan) and the peptides that constitute the same loops in spinach PSII (3JCU, in green). The last modelled amino acids at N- and C- termini of both Arabidopsis loops. Are represented with explicit labels.

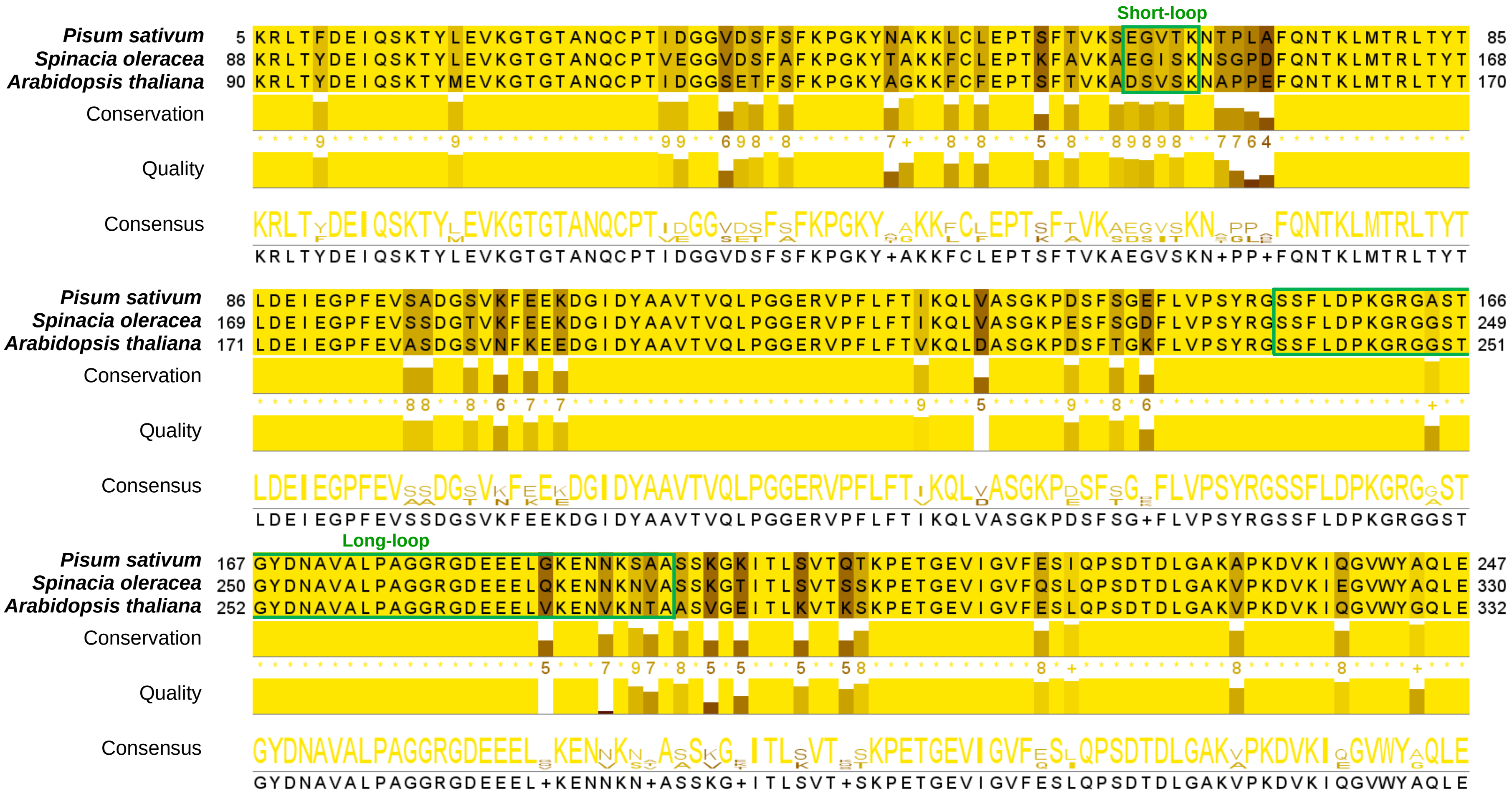

Figure S14 - Sequence alignment of the PsbO proteins of different higher plants.

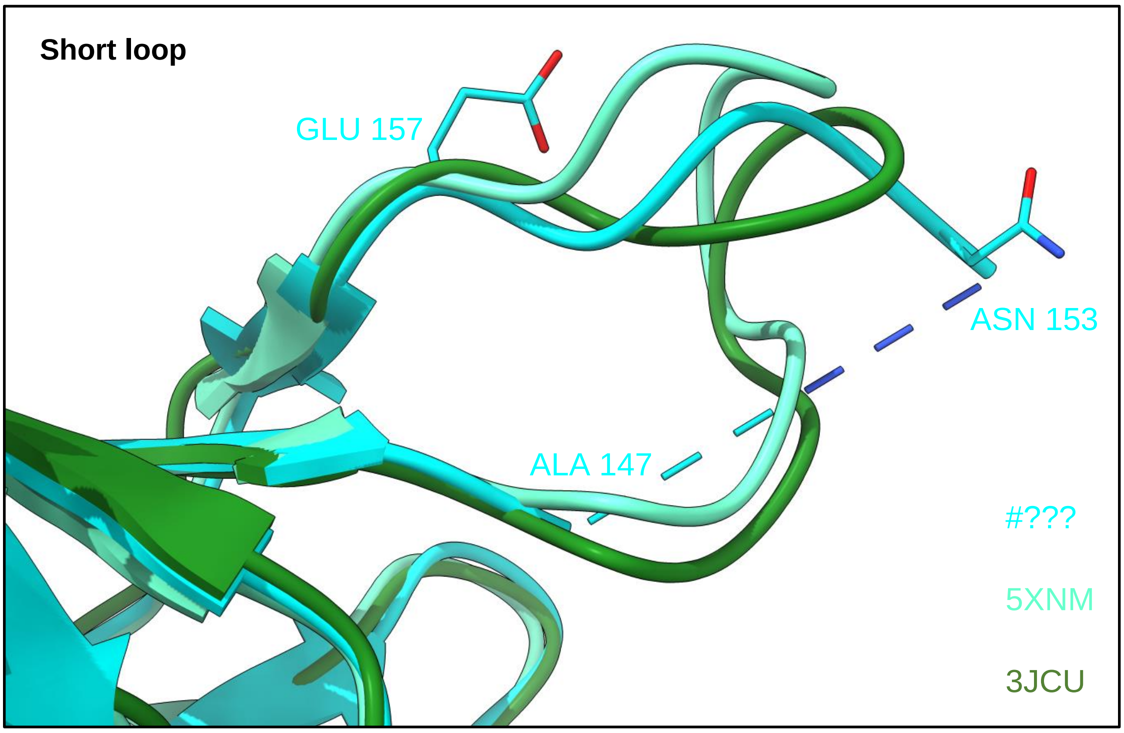


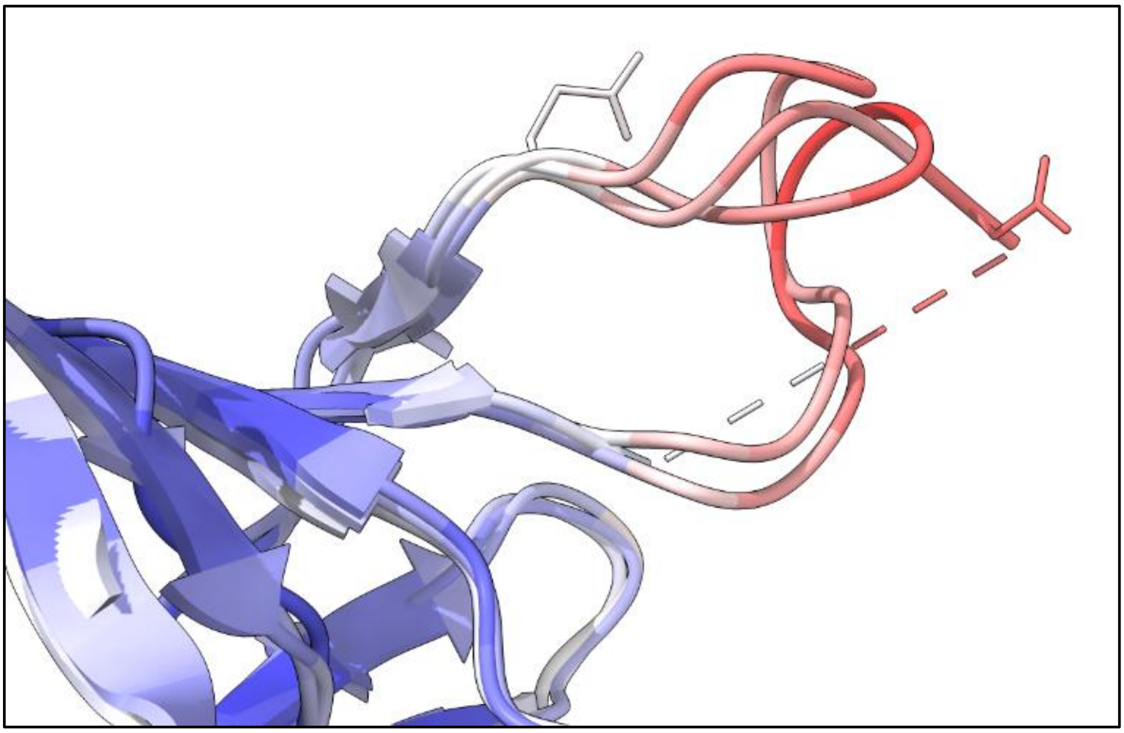

Figure S15 - The PsbO short-loop missing in Arabidopsis PSII model is in a region of several non-conserved amino acids (position 153-157, according to Arabidopsis numbering). In accordance with 5XNM and 3JCU models, which represent the missing amino acids in #???, all the amino acids in the loop region have a high b-factor, factors which likely justify the flexibility and difficulty to model such loop in Arabidopsis.

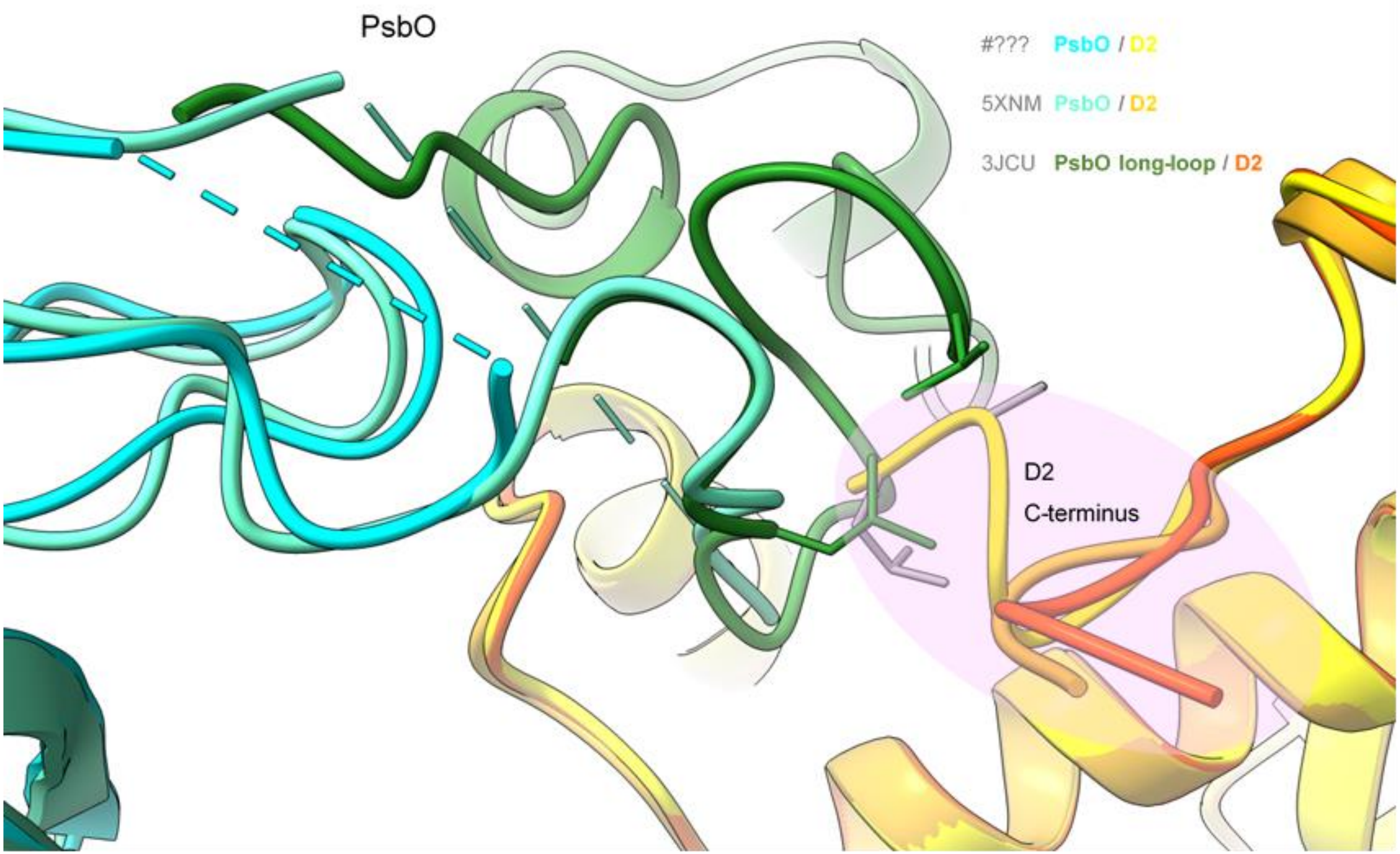


Figure S16 – Representation of the PsbO protein long-loop and the D2 C-terminus of the different higher plants PSII, with a different conformation for Arabidopsis PSII. Such twist occurs at the peptide bond Gly350-Asn351, conflicting with original location of PsbO long-loop.
