## Supplemental Tables for "High-resolution model of Arabidopsis Photosystem II reveals the consequences of digitonin-extraction"

*Table S1 - Statistics of cryo-EM data and structural analysis of the C2S2M2-type PSII supercomplex refined at 2.8 Å resolution. This table is to be completed with the final data after validation upon PDB deposition.*

| | PSII $C_2S_2M_2$ (EMDB-####) (PDB #???) |
|---|---|
| **Data collection and processing** | |
| Hardware | |
| Microscope | Titan Krios |
| Detector (mode) | Gatan K2 BioQuantum (counted) |
| Voltage (keV) | 300 |
| Spherical aberration | 2.7 |
| Magnification | 165 000 × |
| Electron exposure (e/Å$^2$) | 59.7 |
| Defocus range (µm) | -1.5 to -3.0 |
| Pixel size (Å) | 0.82 |
| Symmetry imposed | C2 |
| Initial particle images (no.) | 416 262 |
| Final particle images (no.) | 110 659 |
| Map resolution (Å) | 2.79 ($C_2S_2$ masked) / 3.13 ($C_2S_2M_2$) |
| FSC threshold | 0.143 |
| **Refinement** | |
| Initial models used (PDB code) | 5MDX and 5XNM |
| Model composition | |
| Non-hydrogen atoms | |
| Protein residues | |
| Ligands | |
| B factors (Å$^2$) | |
| Protein | |
| Ligand | |
| R.m.s. deviations | |
| Bond lengths (Å) | |
| Bond angles (°) | |
| Validation | |
| MolProbity score | |
| Clashscore | |
| Poor rotamers (%) | |
| Ramachandran plot | |
| Favored (%) | |
| Allowed (%) | |
| Disallowed (%) | |

*Table S2 - Details on the modelled PSII core subunits. The most common protein names can be found in bold, while alternative names are contained within parenthesis. The molecular weight and pI—computed using ProtParam tool from the Expasy server—regarding the full sequences of mature proteins (according with the respective UniProt entries, accession code preceded by a greater-than symbol before the sequence). The sequences have amino acids letters coloured in 2 distinct colours:* ***black****, all the amino acids modelled in #XXX;* ***red****, amino acids yet unobserved/undistinguishable in a 3D map of* Arabidopsis thaliana*. Modelling completeness is the proportion of modelled in amino acid in #XXX with respect to the mature protein sequence.*

| Chain ID | Protein Name | Gene name | Molecular weight (kDa) | pI | Sequence (mature protein) | Modelling completeness (%) | Model fit into the electrostatic potential map |
|---|---|---|---|---|---|---|---|
| A,a | **D1** ($Q_B$ protein) | PSBA | 37.98 | 5.21 | >P83755<br>TAILERRESESLWGRFCNWITSTENRLYIGWFGVLMIPTLLTATS<br>VFIIAFIAAPPVDIDGIREPVSGSLLYGNNIISGAIIPTSAAIGLHFY<br>PIWEAASVDEWLYNGGPYELIVLHFLLGVACYMGREWELSFRL<br>GMRPWIAVAYSAPVAAATAVFLIYPIGQGSFSDGMPLGISGTF<br>NFMIVFQAEHNILMHPFHMLGVAGVFGGSLFSAMHGSLVTSS<br>LIRETTENESANEGYRFGQEEETYNIVAAHGYFGRLIFQYASFNN<br>SRSLHFFLAAWPVVGIWFTALGISTMAFNLNGFNFNQSVVDS<br>QGRVINTWADIINRANLGMEVMHERNAHNFPLDLA | 98.8 | 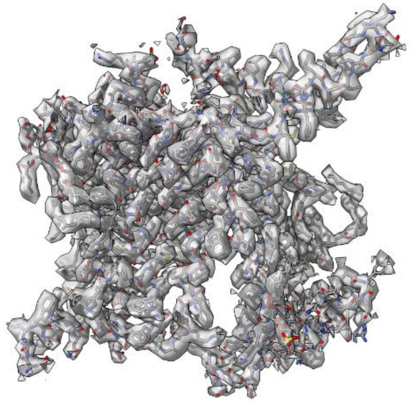 |
| B,b | **CP47** | PSBB | 56.04 | 6.40 | >P56777<br>MGLPWYRVHTVVLNDPGRLLAVHIMHTALVAGWAGSMALY<br>ELAVFDPSDPVLDPMWRQGMFVIPFMTRLGITNSWGGWNIT<br>GGTITNPGLWSYEGVAGAHIVFSGLCFLAAIWHWVYWDLEIFC<br>DERTGKPSLDLPKIFGIHLFLSGVACFGFGAFHVTGLYGPGIWVS<br>DPYGLTGKVQPVNPAWGVEGFDPFVPGGIASHHIAAGTLGILA<br>GLFHLSVRPPQRLYKGLRMGNIETVLSSSIAAVFFAAFVVAGTM<br>WYGSATTPIELFGPTRYQWDQGYFQQEIYRRVSAGLAENQSLS<br>EAWAKIPEKLAFYDYIGNNPAKGGLFRAGSMDNGDGIAVGWL<br>GHPVFRNKEGRELFVRRMPTFFETFPVVLVDGDGIVRADVPFR<br>RAESKYSVEQVGVTVEFYGGELNGVSYSDPATVKKYARRAQLG<br>EIFELDRATLKSDGVFRSSPRGWFTFGHASFALLFFFGHIWHGA<br>RTLFRDVFAGIDPDLDAQVEFGAFQKLGDPTTKRQAV | 95.9 | 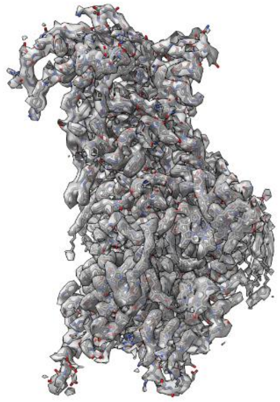 |

| | | | | | | | |
|---|---|---|---|---|---|---|---|
| C,c | **CP43** | PSBC | 50.04 | 6.34 | >P56778<br>TLFNGTLALAGRDQETTGFAWWAGNARLINLSGKLLGAHVAH<br>AGLIVFWAGAMNLFEVAHFVPEKPMYEQGLILLPHLATLGWG<br>VGPGGEVIDTFPYFVSGVLHLISSAVLGFGGIYHALLGPETLEESF<br>PFFGYVWKDRNKMTTILGIHLILLGVGAFLLVFKALYFGGVYDT<br>WAPGGGDVRKITNLTLSPSVIFGYLLKSPFGGEGWIVSVDDLED<br>IIGGHVWLGSICIFGGIWHILTKPFAWARRALVWSGEAYLSYSL<br>AALSVCGFIACCFVWFNNTAYPSEFYGPTGPEASQAQAFTFLV<br>RDQRLGANVGSAQGPTGLGKYLMRSPTGEVIFGGETMRFWD<br>LRAPWLEPLRGPNGLDLSRLKKDIQPWQERRSAEYMTHAPLGS<br>LNSVGGVATEINAVNYVSPRSWLSTSHFVLGFFLFVGHLWHAG<br>RARAAAAGFEKGIDRDFEPVLSMTPLN | 94.3 | 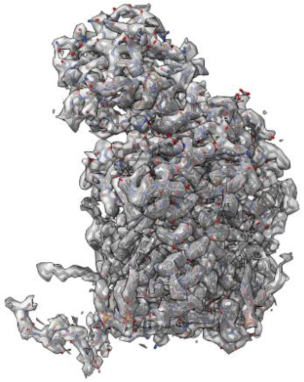 |
| D,d | **D2**<br><br>($Q_A$ protein) | PSBD | 39.42 | 5.46 | >P56761<br>TIALGKFTKDEKDLFDIMDDWLRRDRFVFVGWSGLLLFPCAYF<br>ALGGWFTGTTFVTSWYTHGLASSYLEGCNFLTAAVSTPANSLA<br>HSLLLLWGPEAQGDFTRWCQLGGLWAFVALHGAFALIGFMLR<br>QFELARSVQLRPYNAIAFSGPIAVFVSVFLIYPLGQSGWFFAPSF<br>GVAAIFRFILFFQGFHNWTLNPFHMMGVAGVLGAALLCAIHG<br>ATVENTLFEDGDGANTFRAFNPTQAEETYSMVTANRFWSQIF<br>GVAFSNKRWLHFFMLFVPVTGLWMSALGVVGLALNLRAYDF<br>VSQEIRAAEDPEFETFYTKNILLNEGIRAWMAAQDQPHENLIFP<br>EEVLPRGNAL | 97.2 | 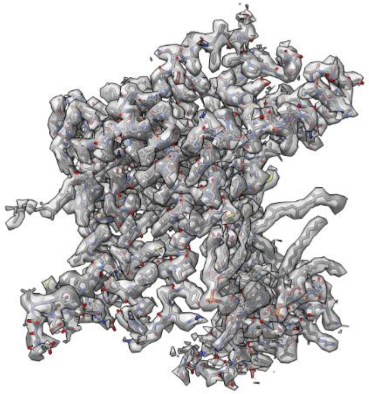 |
| E,e | **Cytochrome $b_{559}\alpha$** | PSBE | 9.39 | 4.83 | >P56779<br>MSGSTGERSFADIITSIRYWVIHSITIPSLFIAGWLFVSTGLAYDV<br>FGSPRPNEYFTESRQGIPLITGRFDSLEQLDEFSRSF | 79.5 | 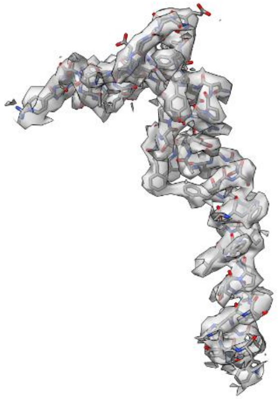 |

| | | | | | | | |
|---|---|---|---|---|---|---|---|
| F,f | **Cytochrom eb$_{559}$β** | PSBF | 4.42 | 10.74 | >P62095<br>MTIDRTYPIFTVRWLAVHGLAVPTVSFLGSISAMQFIQR | 74.4 | 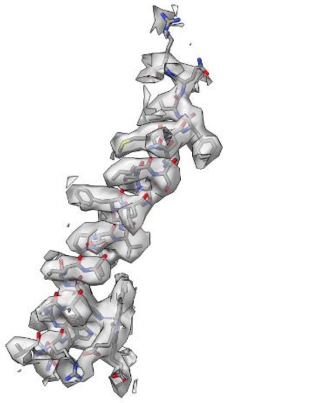 |
| H,h | **Phosphoprotein H** (PSII-H, 10 kDa phosphoprotein) | PSBH | 7.57 | 6.32 | >P56780<br>ATQTVEDSSRSGPRSTTVGKLLKPLNSEYGKVAPGWGTTPLMGVAMALFAVFLSIILEIYNSSVLLDGISVN | 83.3 | 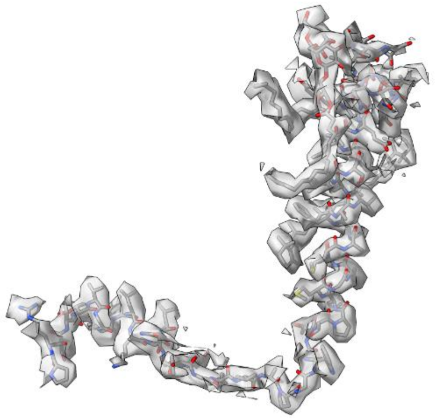 |
| I,i | **Protein I** (PSII-I, 4.8 kDa protein) | PSBI | 4.17 | 5.94 | >P62100<br>MLTLKLFVYTVVIFFVSLFIFGFLSNDPGRNPGREE | 97.2 | 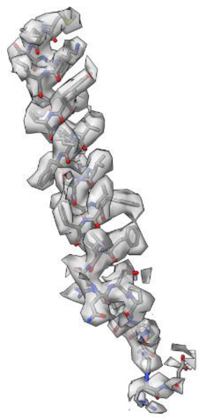 |

| | | | | | | | |
|---|---|---|---|---|---|---|---|
| K,k | **Protein K** (PSII-K) | PSBK | 4.24 | 6.07 | >P56782<br>KLPEAYAFLNPIVDVMPVIPLFFLLLAFVWQAAVSFR | 100.0 | 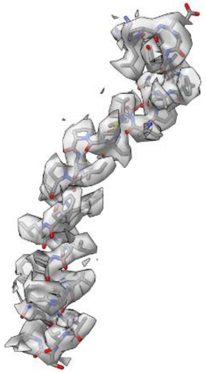 |
| L,l | **Protein L** (PSII-L) | PSBL | 4.47 | 4.53 | >P60129<br>MTQSNPNEQSVELNRTSLYWGLLLIFVLAVLFSNYFFN | 94.7 | 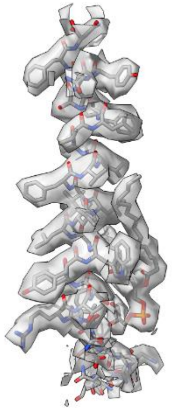 |
| M,m | **Protein M** (PSII-M) | PSBM | 3.78 | 4.37 | >P62109<br>MEVNILAFIATALFILVPTAFLLIIYVKTVSQND | 94.12 | 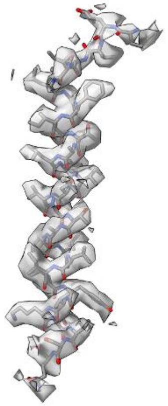 |

| | | | | | | | |
|---|---|---|---|---|---|---|---|
| O,o | **Oxygen-evolving enhancer protein 1-1**, (OEE1, 33 kDa Manganese-stabilizing protein 1, MSP-1, OEC 33 kDa subunit) | PSBO1 | 26.57 | 4.93 | >P23321<br>EGAPKRLTYDEIQSKTYMEVKGTGTANQCPTIDGGSETFSFKPG<br>KYAGKKFCFEPTSFTVKADSVSKNAPPEFQNTKLMTRLTYTLDEI<br>EGPFEVASDGSVNFKEEDGIDYAAVTVQLPGGERVPFLFTVKQL<br>DASGKPDSFTGKFLVPSYRGSSFLDPKGRGGSTGYDNAVALPA<br>GGRGDEEELVKENVKNTAASVGEITLKVTKSKPETGEVIGVFESL<br>QPSDTDLGAKVPKDVKIQGVWYGQLE | 79.8 | 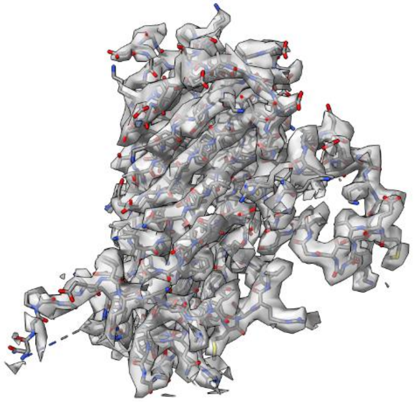 |
| T,t | **Protein Tc** (PSII-T) (Chloroplast T) | PSBTC | 3.82 | 9.52 | >P61839<br>MEALVYTFLLVSTLGIIFFAIFFREPPKISTKK | 87.9 | 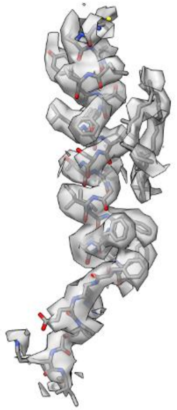 |
| U,u | **Protein Tn** (PSII-T) (nucleus T, Extrinsic T) | PSBTN | 3.17 | 9.70 | >Q39195<br>EPKRGTEAAKKKYAQVCVTMPTAKICRY | 89.3 | 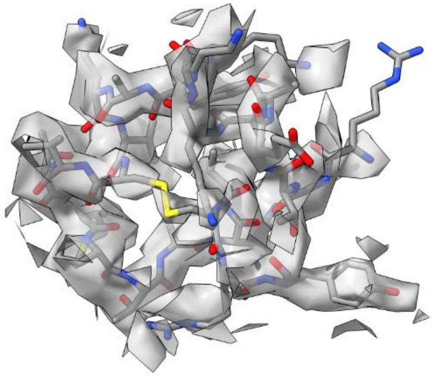 |

| | | | | | | | |
|---|---|---|---|---|---|---|---|
| W,w | **Protein W** (PSII-W, PSII 6.1 kDa protein) | PSBW | 6.04 | 3.67 | >Q39194 LVDERMSTEGTGLPFGLSNNLLGWILFGVFGLIWTFFFVYTSSLE EDEESGLSL | 100.0 | 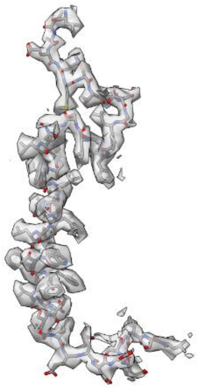 |
| X,x | **Protein X** (PSII-X) | PSBX | 4.18 | 9.99 | >Q9SKI3 AGSGISPSLKNFLLSIASGGLVLTVIIGVVVGVSNFDPVKRT | 85.7 | 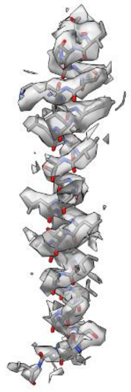 |
| Z,z | **Protein Z** (PSII-Z) | PSBZ, YCF9 | 6.57 | 5.59 | >P56790 MTIAFQLAVFALIITSSILLISVPVVFASPDGWSSNKNVVFSGTSL WIGLVFLVGILNSLIS | 100.0 | 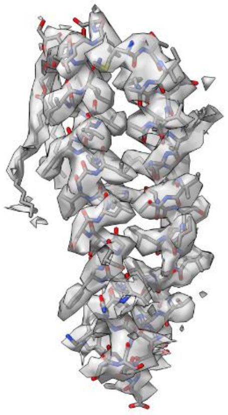 |

*Table S3 - Details on the modelled PSII minor and major antennas. The most common protein names can be found in bold, while alternative names are contained within parenthesis. The molecular weight and pI—computed using ProtParam tool from the Expasy server—regarding the full sequences of mature proteins (according with the respective UniProt entries). The protein sequences presented correspond exclusively to the amino acids present in our model.*

| Chain ID | Protein Name | Gene name | Molecular weight (kDa) | pI | Sequence of modelled protein | Model fit into the electrostatic potential map |
|---|---|---|---|---|---|---|
| G,g | **Chlorophyll a-b binding protein 1**<br><br>(Chlorophyll a-b protein 140, CAB-140, LHCII type I CAB-1)<br><br>**S-Trimer** | LHCB1.3 | 24.86 | 5.12 | >P04778<br>GSPWYGSDRVKYLGPFSGESPSYLTGEFPGDYGWDTAGLSADP<br>ETFARNRELEVIHSRWAMLGALGCVFPELLARNGVKFGEAVW<br>FKAGSQIFSDGGLDYLGNPSLVHAQSILAIWATQVILMGAVEGY<br>RVAGNGPLGEAEDLLYPGGSFDPLGLATDPEAFAELKVKELKNG<br>RLAMFSMFGFFVQAIVTGKGPIENLADHLADPV | 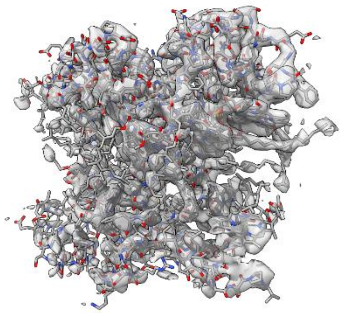 |
| N,n | | | | | >P04778<br>SPWYGSDRVKYLGPFSGESPSYLTGEFPGDYGWDTAGLSADPE<br>TFARNRELEVIHSRWAMLGALGCVFPELLARNGVKFGEAVWF<br>KAGSQIFSDGGLDYLGNPSLVHAQSILAIWATQVILMGAVEGY<br>RVAGNGPLGEAEDLLYPGGSFDPLGLATDPEAFAELKVKELKNG<br>RLAMFSMFGFFVQAIVTGKGPIENLADHLA | 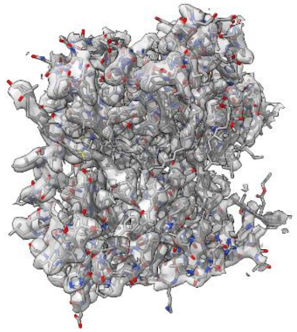 |
| Y,y | | | | | >P04778<br>GSPWYGSDRVKYLGPFSGESPSYLTGEFPGDYGWDTAGLSADP<br>ETFARNRELEVIHSRWAMLGALGCVFPELLARNGVKFGEAVW<br>FKAGSQIFSDGGLDYLGNPSLVHAQSILAIWATQVILMGAVEGY<br>RVAGNGPLGEAEDLLYPGGSFDPLGLATDPEAFAELKVKELKNG<br>RLAMFSMFGFFVQAIVTGKGPIENLADHLADPVNNNAWAF | 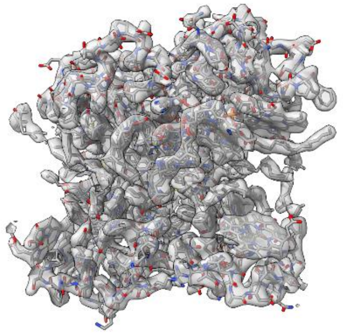 |

| | | | | | | |
|---|---|---|---|---|---|---|
| 1,5 | **Chlorophyll a-b binding protein 1**<br><br>**M-Trimer** | LHCB1.4 | 23.67 | 4.63 | > Q39142<br>SPWYGSDRVKYLGPFSGEPPSYLTGEFPGDYGWDTAGLSADPE<br>TFARNRELEVIHSRWAMLGALGCVFPELLARNGVKFGEAVWF<br>KAGSQIFSDGGLDYLGNPSLVHAQSILAIWATQVILMGAVEGY<br>RVAGDGPLGEAEDLLYPGGSFDPLGLATDPEAFAELKVKELKNG<br>RLAMFSMFGFFVQAIVTGKGPLENLADHLADPVNNNAWAFA<br>TNFVPGK | 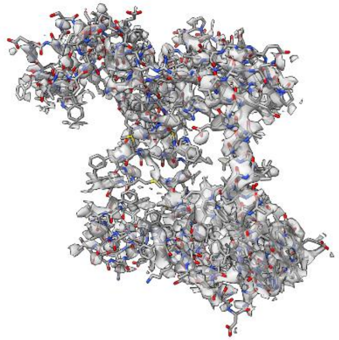 |
| 2,6 | **Chlorophyll a-b binding protein 3**<br>(LHCB3*1)<br><br>**M-Trimer** | LHCB3 | 26.43 | 4.85 | >Q9S7M0<br>ASSFNPLRDVVSLGSPKYTMGNDLWYGPDRVKYLGPFSVQTPS<br>YLTGEFPGDYGWDTAGLSADPEAFAKNRALEVIHGRWAMLG<br>AFGCITPEVLQKWVRVDFKEPVWFKAGSQIFSEGGLDYLGNPN<br>LVHAQSILAVLGFQVILMGLVEGFRINGLDGVGEGNDLYPGGQ<br>YFDPLGLADDPVTFAELKVKEIKNGRLAMFSMFGFFVQAIVTGK<br>GPLENLLDHLDNPVANNAWAFATKFAPGA | 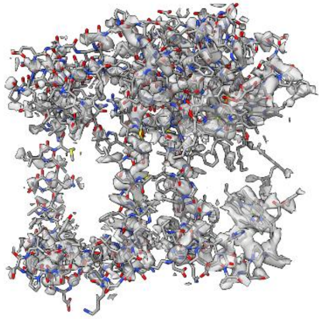 |
| 3,7 | **Chlorophyll a-b binding protein 1**<br><br>**M-Trimer** | LHCB1.4 | 23.67 | 4.63 | > Q39142<br>SPWYGSDRVKYLGPFSGEPPSYLTGEFPGDYGWDTAGLSADPE<br>TFARNRELEVIHSRWAMLGALGCVFPELLARNGVKFGEAVWF<br>KAGSQIFSDGGLDYLGNPSLVHAQSILAIWATQVILMGAVEGY<br>RVAGDGPLGEAEDLLYPGGSFDPLGLATDPEAFAELKVKELKNG<br>RLAMFSMFGFFVQAIVTGKGPLENLADHLADPVNNNAWAFA<br>TNFVPGK | 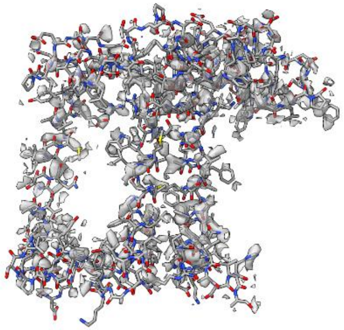 |

| | | | | | | |
|---|---|---|---|---|---|---|
| R,r | Chlorophyll a-b binding protein **CP29**.2 (LHCII protein 4.2) | LHCB4.2 | 28.16 | 5.63 | >Q07473<br>DRPLWYPGAISPDWLDGSLVGDRGFDPFGLGKPAEYLQFDIDS LDQNLAKNLAGDVIGTRTEAADAKSTPFQPYSEVFGIQRFRECE LIHGRWAMLATLGALSVEWLTGVTWQDAGKVELVDGSSYLG QPLPFSISTLIWIEVLVIGYIEFQRNAELDSEKRLYPGGKFFDPLGL AADPEKTAQLQLAEIKHARLAMVAFLGFAVQAAATGKGPLNN WATHLSDPLHTTIIDTFS | 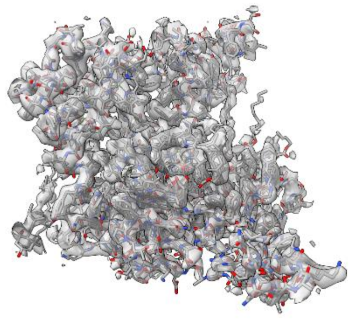 |
| S,s | Chlorophyll a-b binding protein **CP26**, (LHCIIc, Light-harvesting complex II protein 5) | LHCB5 | 25.21 | 4.97 | >Q9XF89<br>DELAKWYGPDRRIFLPDGLLDRSEIPEYLNGEVAGDYGYDPFGL GKKPENFAKYQAFELIHARWAMLGAAGFIIPEALNKYGANCGP EAVWFKTGALLLDGNTLNYFGKNIPINLVLAVVAEVVLLGGAEY YRITNGLDFEDKLHPGGPFDPLGLAKDPEQGALLKVKEIKNGRL AMFAMLGFFIQAYVTGEGPVENLAKHLSDPFGNNLLTVIAG | 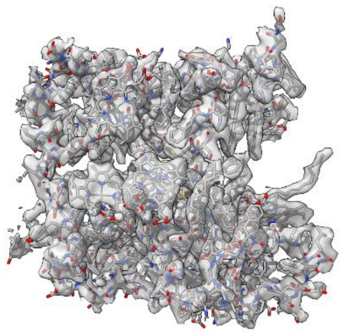 |
| 4,8 | **CP24** | LHCB6 | 23.11 | 5.10 | >Q9LMQ2<br>KKSWIPAVKGGGNFLDPEWLDGSLPGDFGFDPLGLGKDPAFLK WYREAELIHGRWAMAAVLGIFVGQAWSGVAWFEAGAQPDA IAPFSFGSLLGTQLLLMGWVESKRWVDFFNPDSQSVEWATPW SKTAENFANYTGDQGYPGGRFFDPLGLAGKNRDGVYEPDFEKL ERLKLAEIKHSRLAMVAMLIFYFEAGQGKTPLGALG | 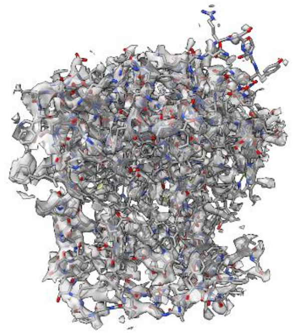 |

*Table S4 – Co-factors present for each higher plant PSII extracted at pH 7.5 (PDB: 5MDX, 5XNM, 3JCU). This table is to be completed with the final data after validation upon PDB deposition.*

| Ligands | | | | | |
|---|---|---|---|---|---|
| **Ligand Name** | **3-letter Code** | **#???** | **5MDX** | **Present in 5XNM** | **Present in 3JCU?** |
| FE (II) ION | FE2 | | A, a | A, a | A, a |
| CHLOROPHYLL A | CLA | | 1, 2, 3, 4, 5, 6, 7, 8, A, B, C, D, G, N, R, S, Y, a, b, c, d, g, n, r, s, y | 1, 2, 3, 4, 5, 6, 7, 8, A, B, C, D, G, N, R, S, Y, a, b, c, d, g, n, r, s, y | A, B, C, D, G, N, R, S, Y, a, b, c, d, g, n, r, s, y |
| PHEOPHYTIN A | PHO | | A, D, a, d | A, a | A, a |
| PROTOPORPHYRIN IX CONTAINING FE | HEM | | E, e | F, f | F, f |
| CHLOROPHYLL B | CHL | | 1, 2, 3, 4, 5, 6, 7, 8, G, N, Y, g, n, y | 1, 2, 3, 4, 5, 6, 7, 8, G, N, Y, g, n, y | G, N, R, S, Y, g, n, r, s, y |
| (3R,3'R,6S)-4,5-DIDEHYDRO-5,6-DIHYDRO-BETA,BETA-CAROTENE-3,3'-DIOL | LUT | | - | 1, 2, 3, 4, 5, 6, 7, 8, G, N, R, S, Y, g, n, r, s, y | G, N, R, S, Y, g, n, r, s, y |
| (3S,5R,6S,3'S,5'R,6'S)-5,6,5',6'-DIEPOXY-5,6,5',6'-TETRAHYDRO-BETA,BETA-CAROTENE-3,3'-DIOL | XAT | | - | 1, 2, 3, 4, 5, 6, 7, 8, G, N, R, Y, g, n, r, y | G, N, R, Y, g, n, r, y |
| (3S,5R,6R,3'S,5'R,6'S)-5',6'-EPOXY-6,7-DIDEHYDRO-5,6,5',6'-TETRAHYDRO-BETA,BETA-CAROTENE-3,5,3'-TRIOL; 9'-CIS-NEOXANTHIN | NEX | | - | 1, 2, 3, 5, 6, 7, G, N, R, S, Y, g, n, r, s, y | G, N, R, S, Y, g, n, r, s, y |
| 1,2-DIPALMITOYL-PHOSPHATIDYL-GLYCEROLE | LHG | | - | 1, 2, 3, 4, 5, 6, 7, 8, B, C, D, G, L, N, R, S, Y, b, c, d, g, l, n, r, s, y | D, G, L, N, R, S, Y, d, g, l, n, r, s, y |
| BETA-CAROTENE | BCR | | - | 4, 8, A, B, C, D, H, T, a, b, c, d, h, t | A, B, C, D, H, a, b, c, d, h |
| CA-MN4-O5 CLUSTER | OEX | - | - | A, a | A, a |

| | | | | | |
|---|---|---|---|---|---|
| 1,2-DI-O-ACYL-3-O-[6-DEOXY-6-SULFO-ALPHA-D-GLUCOPYRANOSYL]-SN-GLYCEROL | SQD | | - | A, B, a, b | A, B, a, b |
| 1,2-DISTEAROYL-MONOGALACTOSYL-DIGLYCERIDE | LMG | | - | A, B, C, D, Z, a, b, c, d, z | A, B, C, D, Z, a, b, c, d, z |
| 2,3-DIMETHYL-5-(3,7,11,15,19,23,27,31,35-NONAMETHYL-2,6,10,14,18,22,26,30,34-HEXATRIACONTANONAENYL-2,5-CYCLOHEXADIENE-1,4-DIONE-2,3-DIMETHYL-5-SOLANESYL-1,4-BENZOQUINONE | PL9 | | - | A, D, a, d | D, d |
| DIGALACTOSYL DIACYL GLYCEROL (DGDG) | DGD | | - | B, C, H, b, c, h | C, H, c, h |
| BICARBONATE ION | BCT | | - | D, d | D, d |
| CHLORIDE ION | CL | | - | - | A,a |
| DIGITONIN | AJP | | - | - | - |
| CALCIUM ION | CA | | - | - | - |

*Table S5 – The different PSII purification conditions and the effects on the OEC environment.*

| | ***Arabidopsis thaliana*** | | ***Pisum sativum*** | | ***Spinacia oleracea*** |
|---|---|---|---|---|---|
| | **#???** | **5MDX** | **5XNM** | **5XNL** | **3JCU** |
| **PsbP?** | NO | NO | NO | YES | YES |
| **PsbQ?** | NO | NO | NO | YES | YES |
| **$Na^+$ as a ligand?** | NO | NO | YES | NO | YES |
| **$Mn_4CaO_5$?** | NO | NO | YES | YES | YES |
| **Starting Membranes** | BBY 1mg/mL washed with 10 mM HEPES-KOH pH [7.2 -7.5] | BBY 1mg/mL washed with 10 mM HEPES-KOH pH 7.5 | Thylakoid membranes 1 mg/mL,<br>in 5 mM MES pH 6.0, 10 mM NaCl, 5 mM $MgCl_2$ and 2 M glycine betaine (MNMβ buffer) | | 500µg grana membranes (BBY?) washed with 1mM EDTA prior to solubilization |
| **Solubilization** | 0.5 mg/mL Chl<br>0.5% (w/v) Digitonin + 0.2% (w/v) Beta-DDM<br>30 minutes | 0.5 mg/mL Chl<br>0.5% (w/v) Digitonin + 0.2% (w/v) Alpha-DDM<br>30 minutes | 2% Alpha-DDM<br>10 mM HEPES pH 7.5 | 2.5% Alpha-DDM<br>10 mM HEPES pH 7.5 | 0.5 mg/mL Chl<br>0.3% Alpha-DDM<br>1 min vortexing<br>10 mM HEPES pH 7.5 |
| **Sucrose Gradient** | Freeze-thawing method.<br>0.35M sucrose gradient<br>10 mM HEPES-KOH pH 7.5<br>0.01% (w/v) digitonin | Freeze-thawing method.<br>0.65M sucrose gradient<br>10 mM HEPES-KOH pH 7.5<br>0.01% (w/v) digitonin | Freeze-thawing method (-80 to 4°C).<br>0.65M sucrose gradient<br>10 mM HEPES pH 7.5 and 0.016% α-DDM | Freeze-thawing method (-80 to 4°C).<br>0.65M sucrose gradient<br>25 mM MES pH 5.7,<br>5 mM $CaCl_2$ and 0.03% α-DDM | Freeze-thawing method (-80 to 4°C).<br>0.65M sucrose gradient<br>10 mM HEPES-KOH pH 7.5<br>0.008% α-DDM |
| **U centrifugation** | 13 000 g for 12 min.<br>41 000 rpm for 7h (200µg Chl) (SW41)<br>Last band was collected | 13 000 g for 10 min.<br>41 000 rpm for 17h (200µg Chl) (SW41) | 247 600 g, 4 °C for 20 h (SW41).<br>3rd band collected | 100 000 g, 4 °C for 15 h (SW41).<br>3rd band collected | 16 000 g for 10 min.<br>247 600g, 4 °C for 21 h (SW41).<br>B9 band Chl a/b ratio around 2.9–3.1 was collected. |
| **Final buffer and pH** | 10mM HEPES-KOH<br>pH 7.5 + 0.01% w/v digitonin | 10mM HEPES-KOH<br>pH 7.5 + 0.01% digitonin | 10mM HEPES<br>pH 7.5 | 25mM MES<br>pH 5.7 | 10mM HEPES<br>pH 7.5 |
| **Final Salt Concentration** | No salt (after BBY prep) | No salt (after BBY prep) | None | 5mM $CaCl_2$ | None |
| **Concentration** | Concentrated to 3.5 mg/mL chl on on Millipore Amicon filter (30 kDa cut-off, 10mL) in rounds of 5.000 g for 30 min and washed in-between w/ 2mL 10mM HEPES-KOH pH 7.5 + 0.01% digitonin. Last concentration step with Vivaspin 500 (100kDa cut-off, 500µL). Final 20µL. | Concentrated to 3.5 mg/mL chl on Millipore Amicon filter (10 kDa cut-off, 10mL) in rounds of 3500 g for 20 min and washed in-between w/ 2mL 10mM HEPES-KOH pH 7.5 + 0.01% digitonin. Final 60µL. | 3 mg/mL (in chlorophyll) | | Concentrated to 3 mg/mL Chl by using a 100-kDa cut-off concentrator. |

*Table S6 – Inter-chlorophyll distances (Mg-Mg) of coupled pairs involved in energy transfer between PSII subunits.*

| Subunit and Chlorophyll ID | Subunit and Chlorophyll ID | Mg-Mg distance (Å) | Location |
|---|---|---|---|
| **M-LHCII$_A$** | **CP29** | | |
| CLA611 | CLA611 | 16.4 | Stroma |
| **CP29** | **CP47** | | |
| CLA603 | CLA610 | 19.2 | Stroma |
| CLA609 | CLA616 | 21.8 | Stroma |
| CLA613 | CLA602 | 23.5 | Lumen |
| **S-LHCII$_A$** | **CP43** | | |
| CLA611 | CLA506 | 16.9 | Stroma |
| CLA612 | CLA506 | 22.1 | Stroma |
| CLA614 | CLA501 | 23.6 | Lumen |
| **S-LHCII$_B$** | **CP26** | | |
| CLA604 | CLA604 | 24.4 | Lumen |
| CLA610 | CLA610 | 26.1 | Stroma |
| **CP26** | **CP43** | | |
| CLA611 | CLA513 | 16.7 | Stroma |
| CLA611 | CLA512 | 17.6 | Stroma |
| CLA614 | CLA503 | 16.7 | Lumen |

| Subunit and Chlorophyll ID | Subunit and Chlorophyll ID | Mg-Mg distance (Å) | Location |
|---|---|---|---|
| **M-LHCII$_A$** | **CP29** | | |
| CLA614 | CHL614 | 14.8 | Lumen |
| **S-LHCII$_A$** | **CP29** | | |
| CHL605 | CHL606 | 18.4 | Lumen |
| CHL605 | CLA604 | 17.9 | Lumen |
| **S-LHCII$_B$** | **CP26** | | |
| CHL605 | CLA604 | 19.2 | Lumen |
| CHL608 | CLA610 | 20.9 | Stroma |
| CHL608 | CLA612 | 21.0 | Stroma |
| **CP26** | **CP43** | | |
| CHL601 | CLA513 | 13.0 | Stroma |
